# Single cell decoding of drug induced transcriptomic reprogramming in triple negative breast cancers

**DOI:** 10.1101/2023.09.19.558329

**Authors:** Farhia Kabeer, Hoa Tran, Mirela Andronescu, Gurdeep Singh, Hakwoo Lee, Sohrab Salehi, Justina Biele, Jazmine Brimhall, David Gee, Viviana Cerda, Ciara O’Flanagan, Teresa Algara, Takako Kono, Sean Beatty, Elena Zaikova, Daniel Lai, Eric Lee, Richard Moore, Andrew J. Mungall, IMAXT Consortium, Marc J. Williams, Andrew Roth, Kieran R. Campbell, Sohrab P. Shah, Samuel Aparicio

## Abstract

**Background:** The encoding of cell intrinsic resistance states in breast cancer reflects the contributions of genomic and non-genomic variation. However, identifying the potential contributions of each requires accurate measurement and subtraction of the contribution of clonal fitness from co-measurement of transcriptional states. Somatic genomic variation in gene dosage, copy number variation, is the dominant mutational mechanism in breast cancer contributing to transcriptional variation and has recently been shown to contribute to platinum chemotherapy resistance states. Here we deploy time series measurements of triple negative breast cancer single cell transcriptomes in conjunction with co-measured single cell copy number associated clonal fitness to identify the contributions of genomic and non-genomic mechanisms to drug associated transcription states.

**Results:** We generated serial scRNA-seq data (126,556 cells) from triple negative breast cancer (TNBC) patient-derived xenograft (PDX) experiments over 2.5 years in duration, and matched it against genomic copy number single cell data from the same biological samples. We show that the cell memory of transcriptional states of TNBC tumors serially exposed to platinum identifies distinct clonal responses within individual tumours. Copy-number clones with high drug fitness leading to clonal sweeps exhibit less transcriptional reversion, whereas clones with weak drug fitness exhibit highly dynamic transcription on drug withdrawal. Pathway analysis shows that copy number associated and copy number independent transcripts converge on epithelial-mesenchymal transition (EMT) and cytokine signaling states associated with resistance. We show from trajectory analysis that transcriptional reversion exhibits hysteresis, indicating that new intermediate transcriptional states are generated by platinum exposure.

**Conclusions:** We discovered that copy number clones with strong genotype associated fitness under platinum became fixed in their states, resulting in minimal transcriptional reversion on drug withdrawal. In contrast clones with weaker fitness undergo non-genomic transcriptional plasticity and these distinct responses co-exist within single tumours. Together the data suggest that copy number associated and copy number independent transcriptional states may contribute to platinum drug resistance within individual tumours. The dominance of genomic or non-genomic mechanisms within individual polyclonal tumours has implications for approaches to restoration of drug sensitivity and re-treatment strategies.

**Data availability:** Uploaded Data URL: https://ega-archive.org/studies/EGAS00001007242

Github manuscript: https://github.com/molonc/drug_resistant_material/

## Background

The concept of gene expression plasticity and evolutionary fixation is fundamental to understanding how cell populations adapt to rapid environmental changes in a growing tumor. As gene expression includes a significant stochastic component, resulting in cell-to-cell variations in mRNA, the sources of this variability include fluctuations in the expression of individual genes and even genetically identical cells can be very different [1–7]. Studies in cell lines and model systems have emphasized the role of non-genomic transcriptional plasticity, either through fixation of stochastic transcriptional states and/or modulation of transcription through epigenetic regulation of transcription. Drug mechanisms and the duration and speed of onset of drug action likely influence the contributions of genomic and non-genomic encoding of transcriptional states. This is emphasized in the discoveries of rare drug tolerant cells and persister cells in the cell populations of untreated cancers [5,6,8,9]. In most instances these approaches have not observed genomic differences, potentially reflecting the speed of action and drug mechanisms studied. However not all studies have systematically sequenced single cells, especially to detect somatic gene dosage variants. Copy number structural variants have greater potential to shape the fitness landscapes of resistance in cancers with genomic instability, because single mutations can affect the gene dosage and transcription levels of potentially hundreds of genes [10,11], creating a potentially large fitness landscape for drug selection. During the evolution of genomically unstable tumours, copy number differences resulting in gene dosage differences between clones may have an extensive effect on transcription and contribute to heritable proportional gene expression differences between clones [10–13]. In triple negative breast cancer (TNBC), gene copy number changes are very abundant and have a major influence on the expression landscape [10,13–15] and are associated with the clinical biology of these cancers.

Few studies have been structured to separate the genome effects of somatic copy number variants from non-genomic transcriptional plasticity. A recent analysis of a different process, colorectal cancer metastasis, has emphasized that in that cancer type, there is only minor variation in genotypes between primary and metastatic colorectal tumours despite ongoing background mutation, implying the primary tumour landscape is under strong selection for stability. In this case, non-genomic transcriptional variation accounts for the majority of the phenotype differences [16,17]. The situation in breast cancers is different, where much greater variation in copy number genotypes has been observed among metastases. Bulk WGS methods can usually only detect differences once clonal selection and fixation has occurred and are thus not well suited to analyzing the dynamics of clonal and non-clonal selection. Thus analysis of the potential contributions of gene dosage mutations and non-genomic transcriptional variation in response to chemotherapy requires single cell analysis of serially sampled populations. Using a scaled single cell genome sequencing technology and population genetic fitness models, we have recently shown that copy number genotype differences among subclones of TNBC can be associated with platinum resistance [18]. Here we have exploited this observation to dissect the dynamics of clone associated and clone independent transcription during the onset and reversion of platinum drug resistance sampling 126,556 single cell transcriptomes of serially platinum treated TNBC patient derived xenografts over a 2.5 years interval. We show that even within a tumour, different subclones may exhibit quite different transcriptional dynamics to drug treatment and withdrawal, reflecting the degree of clonal fixation.

## Results

### Copy number variation shapes PDX single cell transcriptomes in proportion to genomic instability

We set out to measure copy number aberration (CNA) associated and copy number independent transcriptional variation in subpopulations of patient derived xenografts passaged over months to years under neutral (no drug intervention) conditions (Figure 1) and with parallel cisplatin treatment and cisplatin holiday conditions. We established a new dataset of joint tumour cell population measurements utilizing DLP+ single cell genome sequencing (scWGS) to measure clonal structure determined by CNA and scRNA-seq sampled from single cells in the same population from 6 triple negative breast cancer patient-derived xenograft series (Figure 1), including 3 patients where serial platinum treatment samples were obtained. From 126,566 single cell transcriptomes, we captured 55,025 high quality transcriptomes (post filtering), representing 29,475 cells from untreated tumors (Pt1-6), 14,100 cells from cisplatin treated tumors (Pt4,5,6), and 11,450 cells from cisplatin drug holiday tumors (Pt4,5,6, Table 1, Supplementary Table 1).

**Figure 1.**
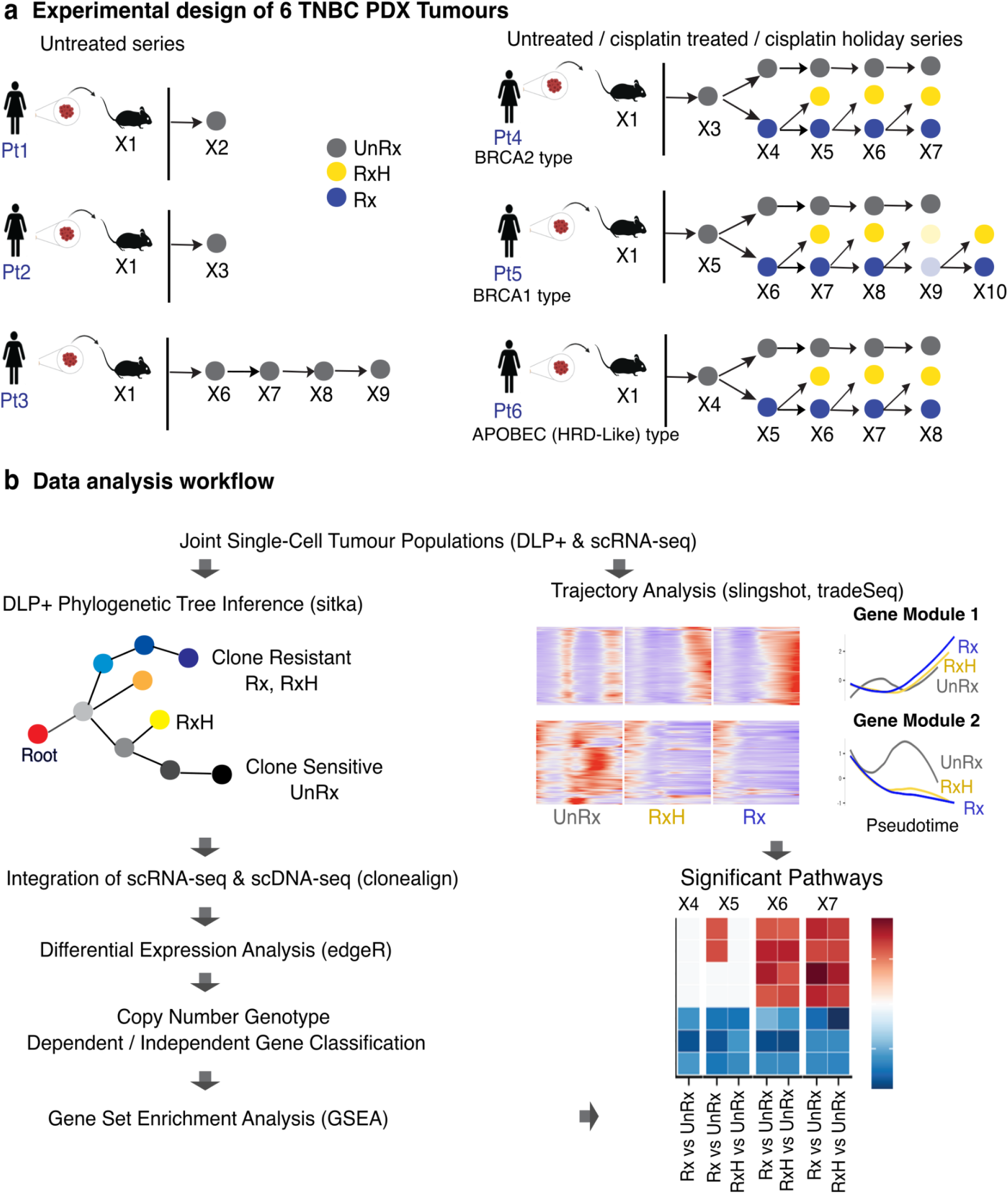
Workflow of experimental and study design for tracking drug induced transcriptome reprogramming. (a) Tumor biopsies from 6 TNBC patients were transplanted in immunodeficient mice. Three TNBC untreated time-series (Pt1-3) and three cisplatin treated time-series (Pt4-6) with its counter drug holiday samples. UnRx: untreated, Rx: cisplatin treated, RxH: cisplatin drug holiday. The treated, drug holiday samples at X9, Pt5 were excluded from analysis due to its sample qualities. (b) Data analysis workflow, including phylogenetic tree inference using DLP single cell copy number profiles from previously published work [19], followed by clonal alignment from DLP copy number to RNA-seq gene expression with clonealign [20]. Differentially expressed genes are then classified into in cis and in trans based on overlapping of bin genomic and gene genomic regions, and based on the positive or negative directions of copy number tendency and gene expression trend (up-regulated or down-regulated). Significant pathways are determined by applying gene set enrichment analysis. Pseudotime analysis highlights the achieved genes with significant change in expression along the trajectories of evolution through drug treatment.

**Table 1.**
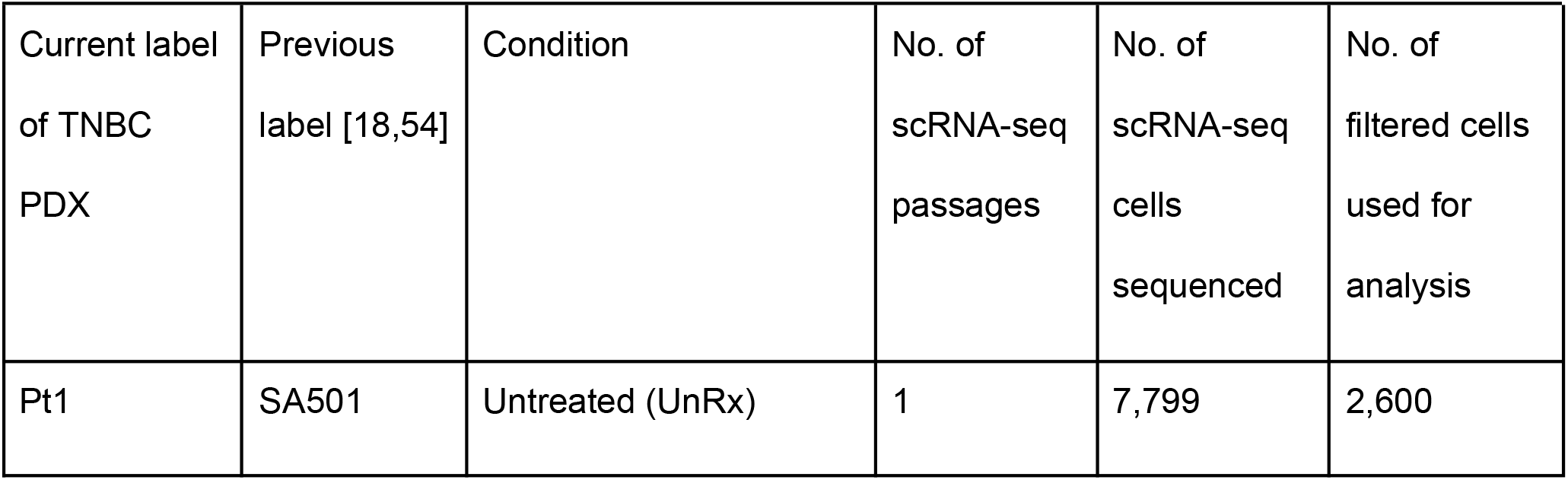
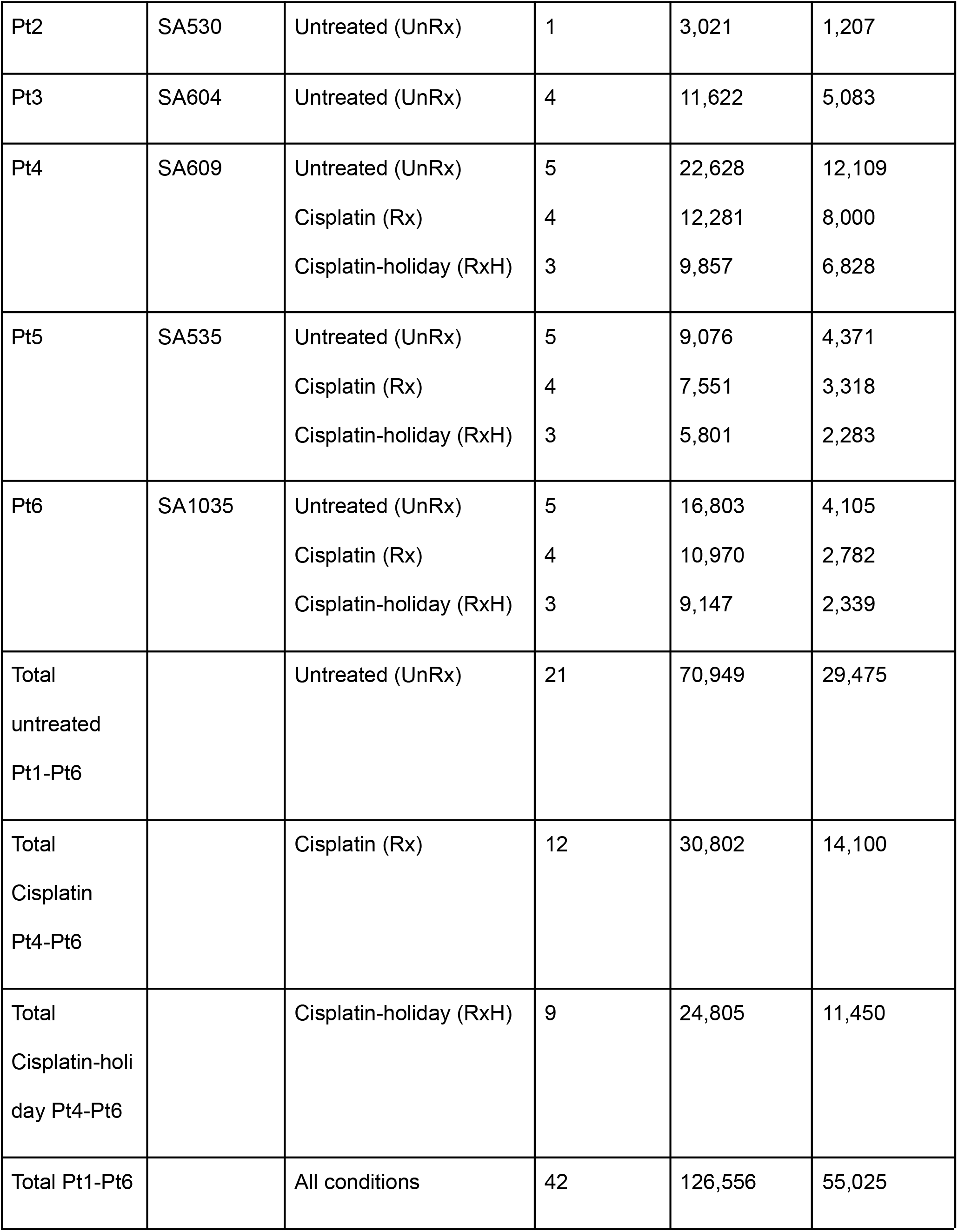
Types and numbers of single cell RNA-seq data generated for this study. See also Supplementary Table 1 for details of each sample.

To assign scRNA-seq transcriptomes to clones, we first generated CNA phylogenetic trees with a Bayesian [19] method from DLP+ single cell genome sequencing of 6 patient PDX lines, to determine the genome fraction occupied by TNBC breast cancer clones. For each patient series we identified CNA clonal populations from major clades (Figure 2a), and copy number profiles for each clone (Figure 2b,c, Supplementary Figures 2-5). Reflecting the background genomic diversity of breast cancer patients [10], the number of major clones varied from 4 (Pt2) to 11 (Pt3,6), while the proportion of genome altered by CNA (Manhattan distance) was <0.06 average CNA distance for Pt2 and Pt6, and > 0.2 for Pt3,4,5, reflecting the natural variation of CNA clonal composition of TNBC [18]. Among three patient lines treated serially with platinum (Pt4,5,6, Figure 1), Pt4 exemplifies a low complexity (6 clones) and intermediate fraction of altered genome (mean=0.11, sd=0.07), Pt6 a high clone complexity (11 clones) and low fraction of altered genome (mean=0.04, sd=0.02), whereas Pt5 exhibited high clonal complexity (10 clones) and high altered genome fraction (mean=0.3, sd=0.16), Figure 2a,b.

**Figure 2.**
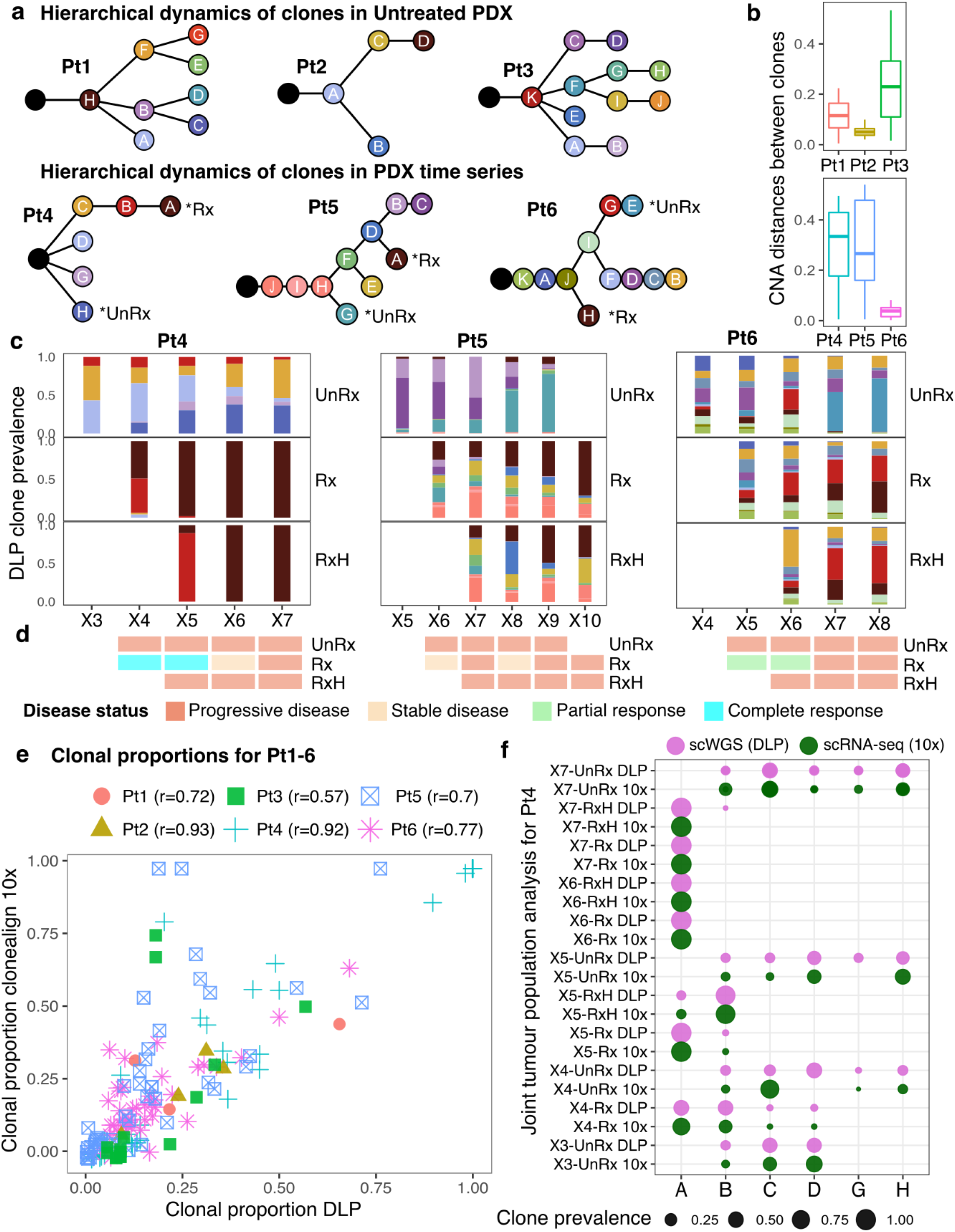
Assignment of structural-copy number clone structure to scRNA-seq. Summary results of phylogenetic trees and clone alignment correlations. (a) Inferred phylogenetic trees of three untreated PDX patients Pt1-3, and three drug treatment time-series data Pt4-6. Resistant, sensitive cell clones were developed at different branches far from each other. (*) denote high fitness coefficient clones in each branch based on the previously published work [18]. (b) Manhattan copy number distances between median copy number profiles of paired clones in each series denote the variance level in the copy number DLP tree. (c) Clonal evolution of cells with/without drug treatment across time is well captured thanks to time-series tracking. (d) Disease classification at each time point and treatment status for Pt4-6. (e) Summary clone alignment Pearson correlation between DLP copy number and RNA-seq gene expression from 6 patients in this study using clonealign package [20]. (f) Pearson correlation between clonal proportions in DLP, and inferred clonal proportions in 10x - scRNA-seq expression at the same passages from patient Pt4 using clonealign. Strong positive correlation demonstrates that clonealign is able to provide accurate alignment.

We have previously measured the clonal fitness of Pt4,5,6 under different passaging conditions [18]. Briefly, the fitness coefficient denotes the relative growth potential rate (increased or attenuated) of a given clone over time and was quantified using a Wright-Fisher inspired Bayesian probabilistic framework named *fitClone*. The fitness coefficient results from this previous analysis for patients Pt4-6 are included in Supplementary Figure 3b, Supplementary Figure 4b, Supplementary Figure 5b and Supplementary Table 2. In this study, clones from the treatment series with high fitness coefficient were labeled as *Rx (resistant to drug), and clones from the untreated series with high fitness coefficient were labeled as *UnRx (sensitive to drug, Figure 2a, Supplementary Table 2). Using mRECIST criteria, ***complete response, partial response*** <50% tumor shrinkage, ***stable disease*** 0% tumor shrinkage or ***progressive disease*** (Figure 2d and Extended Data Fig. 3 of ref. [18]), we observed initial physical tumour responses with onset of resistance within 2 cycles, indicating that early clonal responses were also mirrored by physical tumour volume response.

Next, we mapped scRNA-seq from cells drawn from the same tumour cell populations to the CNA clones defined above, using *clonealign* [20], which probabilistically assigns CNA clone identity to scRNA-seq transcriptomes, assuming that an increase in the copy number of a gene will result in a corresponding increase in that gene’s expression and vice versa (Methods, Supplementary Table 3). The inferred clonal prevalences (Figure 2e,f) of cells in scRNA-seq populations were highly concordant with the clonal prevalences of cells in DLP+ populations (Figure 2c), Pearson correlation coefficient r>0.7 for all but 1 patient, overall correlation r=0.8, p-value<0.05 for all patients, average correlation in three treatment series was greater when cells were under treatment r=0.8+-0.11, p-value<0.05.

Next, having established CNA clone identity of transcriptomes, we investigated differential transcription among clones that had the highest fitness coefficient [18], and/or were the most abundant at each time series sample (see Figure 3a), using *edgeR*, which implements a Negative Binomial model to determine genes whose expression changes between conditions [21] (Methods). The gene expression change strongly correlated with the copy number change (Pearson correlation coefficient > 0.95, Supplementary Figure 6), which is expected given *clonealign*’s model of gene expression – copy number correlation, and consistent with previous findings across many tumour types including breast cancer[10,11].

**Figure 3.**
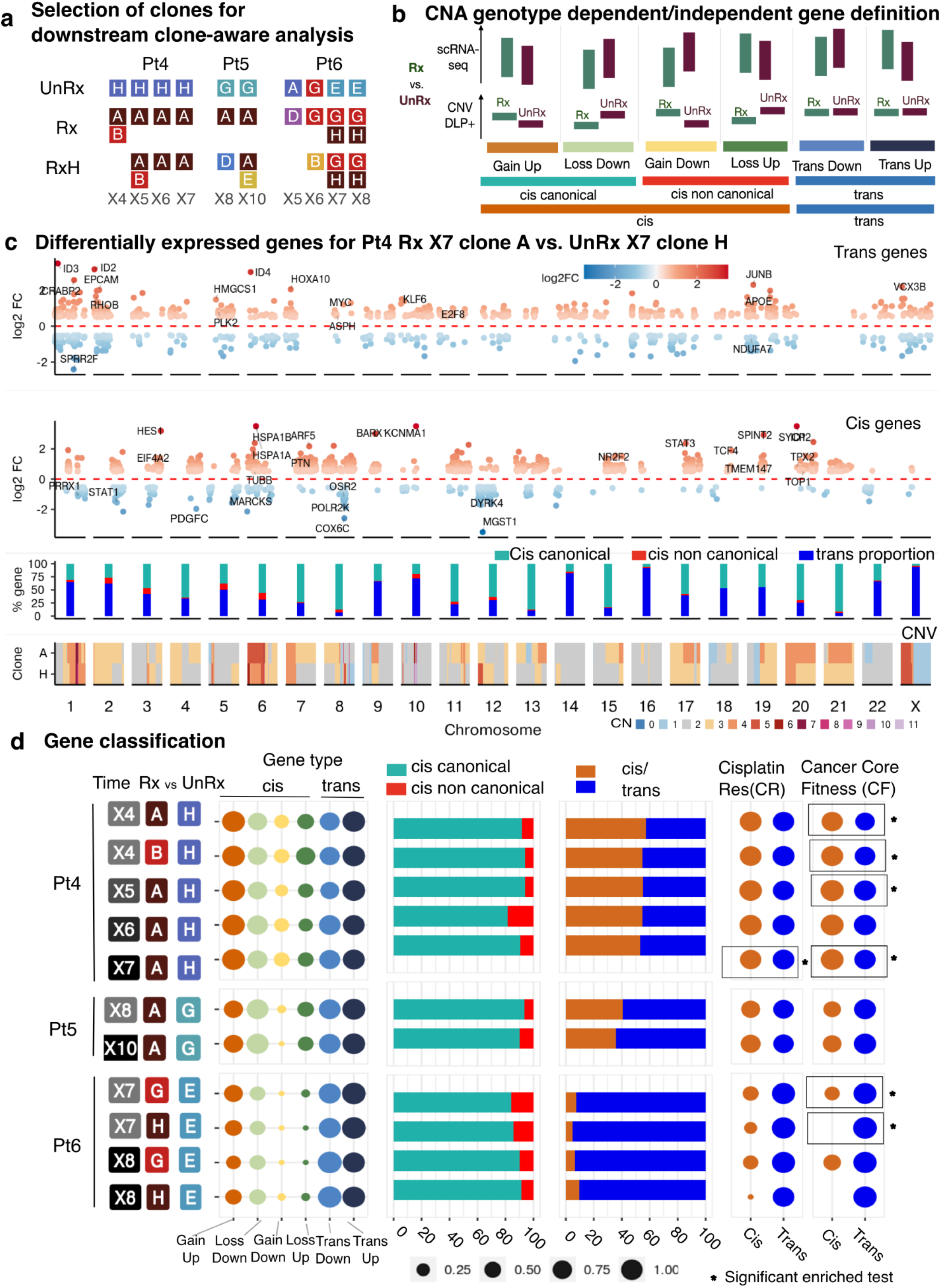
Cis and trans gene proportions and gene set memberships. (a) For Pt4-6, the clones that were the fittest or most abundant and had at least 100 scRNA-seq cells were selected for further clone-aware analysis. (b) Gene classification: genes are divided into cis - CNA genotype dependent genes, and trans - CNA genotype independent genes based on whether the copy number values changed at the overlapping genomic regions of the pair clones comparisons. Further, cis genes are divided into copy-number correlated and non copy-number correlated based on the same or opposite directions of gene expression and copy number values. Genes are then classified based on the direction of change in gene expression (up- or down-regulated) as compared to the direction of change in copy number (gain or loss). (c) Differentially expressed genes for Pt4 between resistant clone A in Rx passage X7 versus sensitive clone H in UnRx passage X7, classified into “trans” genes (CNA genotype independent gene, top panel) and “cis” genes (CNA genotype dependent gene, second panel). Red and blue gradient colors denote the degree of log2 FC in positive and negative directions. Each dot is one significant DE gene with selected condition abs(log2FC)>0.5, FDR<0.01, p value<0.05. Third panel: % of cis copy-number correlated - light green, non copy-number correlated - red, and trans genes - blue color per chromosome. Bottom panel: median copy number profile of genomic regions for the two clones from DLP+ results at corresponding regions. (d) Differentially expressed genes between resistant clones versus sensitive clones in Pt4-6 were classified into different gene types. The dot size represents the proportion of each gene type based on the definition in (b). Gene set membership: mapping of cis and trans genes to two reference gene sets: COSMIC core fitness census genes (CF) [55] and Cisplatin Resistance set (CR) - curated genes list from the latest literature with cisplatin resistance. Results of statistical tests using GSEA [56] gene set enrichment analysis with p-adj values < 0.05, where rectangles with * show significant enrichment of reference gene sets.

Transcripts encoded in regions of clone to clone copy number differences were labelled as *in-cis*, whereas transcripts encoded outside of clonal CNA differences were labelled as *in-trans* (Figure 3b). *In-trans* regions exhibited a larger proportion of up than down regulation differences across treated and untreated tumour (39.5% sd=8.5% of the up-regulated trans genes on average across patients Pt4,5,6 as compared to 26.9% sd=15.9% down-regulated trans, KS test p-value=0.03, Figure 3c, top panel). Regions identified as *in-cis* exhibited the expected (copy-number correlated) directional tendency of differential expression (example Pt4, Figure 3c, lower track), with a small proportion of copy-number anti-correlated *in-cis* transcripts regulated against the direction of clonal copy number difference (example Pt4, Figure 3c, second track). Copy-number correlated and anti-correlated *in-cis* differential expression was evident over all sizes of CNA from sub-chromosomal regions (e.g., Pt4, Chr1, Chr18, Figure 3c, second track) to whole chromosomes (Pt4 Chr 7, Figure 3c, second track, Pt5,6, Supplementary Figure 7).

We identified copy-number correlated and copy-number anti-correlated differential expression and quantified the proportionality of *in-cis* and *in-trans* DE transcripts across the selected clones (Figure 3b,d, Supplementary Figure 8, Supplementary Table 4 and Supplementary Table 5). The fraction of CNA genotype associated in *cis* transcripts varied from 5.1 - 57.4% of the total number of DE genes in Pt4,5,6, reflecting the variance in the size and number of genomic copy number differences (Figure 2b, Supplementary Figures 3a,4a,5a) between clones for each patient. The proportion of *in-trans* transcripts up- and down-regulated between pairs of clones was similar in all untreated patients (Supplementary Figure 8). However the DE variance of *in-trans* transcripts was greater (*in-trans* up-regulated, sd=4%, down-regulated sd=3.05%) than *in-cis* region transcript (*in-cis* up regulated sd=0.49%, *in-cis* down sd=1.22%; Levene’s test, p-value<0.05 for up-regulated genes logFC values between cis, trans genes in Pt4, Pt5, and for down-regulated genes in Pt4, Pt6). In contrast, a small proportion of *in-cis* transcripts (3.1%, sd=2.2%, Figure 3d) in each of the Pt4,5,6 patient treatment series exhibited *in-cis* copy-number anti-correlated differential expression, indicating a relatively constant fraction of transcripts escaping gene segment copy number dosage effects. This copy-number anti-correlated DE transcript proportionality was stable across time points (Pt4 sd=2.8%, Pt5 sd=0.7%, Pt6 sd=0.2%). Copy-number anti-correlated transcripts have been noted in several studies [10,11,22]. Copy-number anti-correlated transcripts may arise from small CNAs that were missed (for example the PDGFC gene, see Figure 3c, second panel, chromosome 4), or by mis-calling the locations of CNA segment boundaries (on average 11.7% of the copy-number anti-correlated transcripts overlap with the copy number bin boundaries: 11.6% for Pt4, 13.4% for Pt5 and 10.6% for Pt6). However, multiple genes that span the same CN segment are likely to be true copy-number anti-correlated genes (see Figure 3c, second panel, chromosome 2), suggesting that a small proportion of transcripts are biologically counter regulated.

We asked whether DE transcripts reflected possible resistance mechanisms. Through GSEA analysis using curated cisplatin resistance genes (Supplementary Table 6) and published cancer gene sets [23], we found several clones that exhibited statistically significant enrichment of differentially expressed core fitness genes, whereas only one clone pair exhibited enrichment of known cisplatin resistance genes (Figure 3d, Supplementary Table 6), indicating that additional resistance pathways remain to be discovered.

### Clonal and non-clonal transcriptional responses to therapy reveal intratumoural heterogeneity

Having mapped *cis (CNA-clonal)* and *trans* region single cell transcription, we examined the transcription modulation associated with cisplatin treatment and with treatment withdrawal from in-parallel treatment holiday transplants of previously drug exposed tumours [18]. This partially mimics cyclical dosing of patients with platinum.

Investigating the scRNA-seq transcripts that are stable between treatment and holiday transcriptomes, but expressed differentially from the untreated state (Figure 4a,b), we observed that in all patients drug treatment induces more transcripts than are repressed, (Figure 4a-d and Supplementary Figure 9b, Supplementary Table 7), ranging from ∼250-2000 per comparison. The proportion of genes exhibiting increased (induced) stable drug expression (eg. RAC3, Pt4, Figure 4d) was higher (66%) than stably decreased (repressed) transcripts (e.g. DICER1 Pt6, Figure 4d) in all series except for the last time points of Pt5 and Pt6 (43% and 35%, respectively). The proportions of in-cis versus in-trans genes that contribute to the stable expression state of drug exposed cells (36-56% for Pt4, 10-53% for Pt5 and 0-12% for Pt6, Figure 4b) were generally higher than those of untreated passages (Figure 3d). In accordance with our prior observations [18] of copy-clone linked platinum fitness, the distance in copy number space (number of amplification events, depletion events) and copy-clone fitness coefficient values (1+s) are higher in Pt4 compared with Pt5, Pt6.

**Figure 4.**
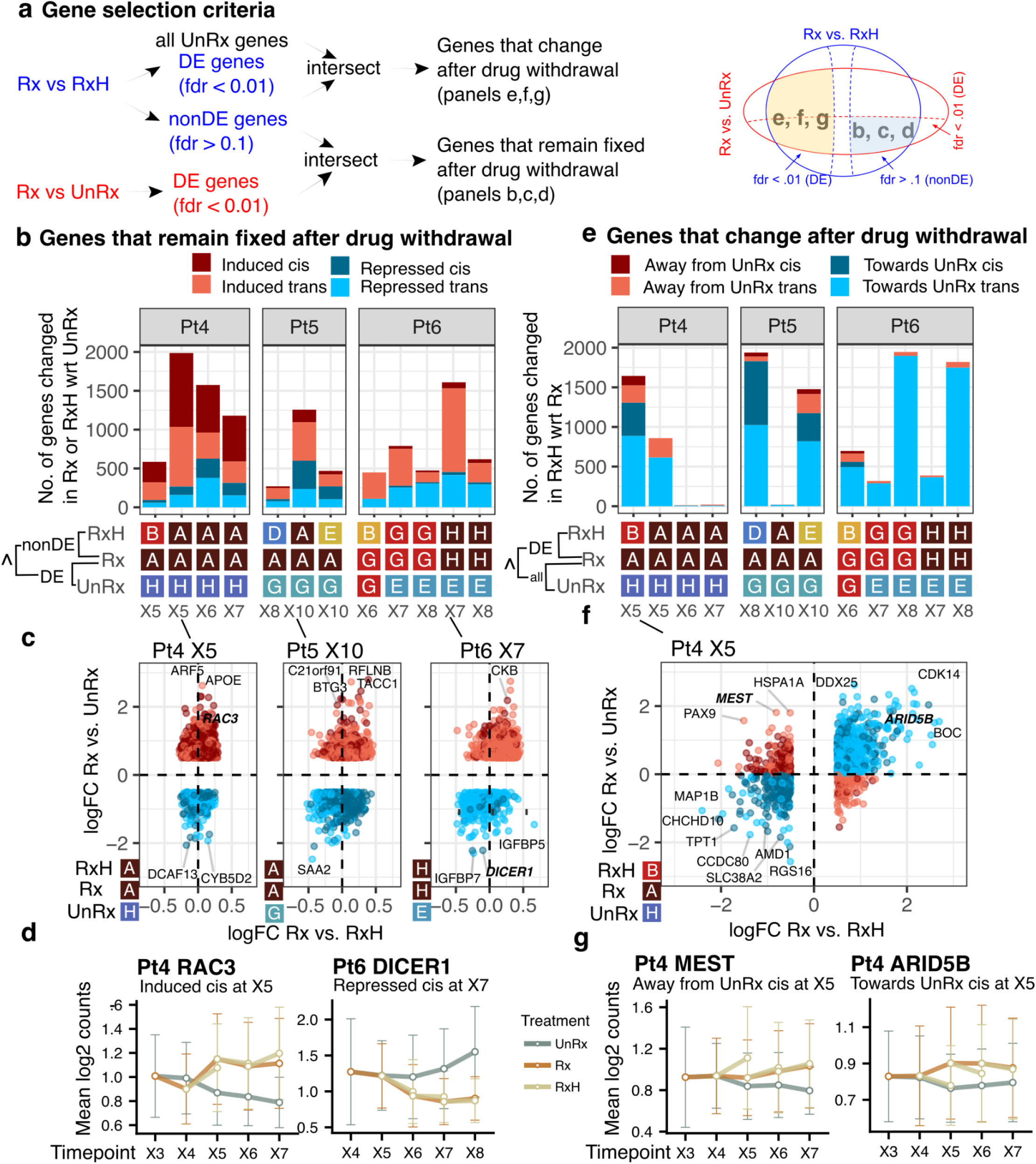
Gene level dynamic changes. (a) Schematic showing our gene selection criteria for Pt4-6: genes in (b,c,d) are differentially expressed (DE, fdr < 0.01, |log2 fold change| = |logFC| > 0.5) at Rx vs. UnRx and non-DE (fdr > 0.1) at Rx vs. RxH; genes in (e,f,g) are DE at Rx vs. RxH and intersected with all the genes in Rx and UnRx. The clones in Figure 3a were used for comparisons. (b) Number of cis or trans treatment induced (logFC>0.5) and repressed (logFC<-0.5) genes for Pt4-6, when comparing the clones listed in the bottom panel. Genes are selected as explained in panel a. The cis and trans gene annotation uses the copy numbers at Rx and UnRx. (c) Scatter plots for logFC of Rx vs. UnRx (y axis) against logFC of Rx vs. UnRx (x axis) for the comparisons with the largest numbers in panel b (second stacked bar for P4 and Pt5 and fourth stacked bar for Pt6). Each point is a gene. The color legend is the same as in panel b. (d) Two examples of induced and repressed genes for Pt4 and Pt6. (e) Number of cis or trans holiday diverged genes, split in two main categories: “towards UnRx” if logFC of Rx vs. RxH has the same sign as logFC of Rx vs. UnRx, and “away from UnRx” otherwise. The comparisons are as explained in panel a. The cis and trans gene annotation uses the copy numbers at Rx and RxH. (f) Scatter plots for logFC of Rx vs. UnRx (y axis) against logFC of Rx vs. UnRx (x axis) for Pt4 X5 (UnRx:H, Rx: A, RxH:B). Each point is a gene. The color legend is the same as in panel e. (g) Two examples of diverged genes away from (left) and towards (right) UnRx.

We asked how transcription changed over time in the same clones, where these could be observed in sufficient abundance over multiple passages. We have previously shown that Pt4 exhibited the strong genotypic clonal selection for platinum resistance, with eventual domination (by passage X6,7, Figure 2c) of a single resistant clone labeled A. In Pt4 comparisons of the untreated high fitness clone H, which is in a distant branch to strongly platinum resistant clone A (Figure 2a), suggested a gradual decline in differential expression over time (Figure 4b). However, differential expression with untreated clone C, which is in the same clade as clone A, showed fewer transcriptional differences (Supplementary Figure 9a). In Pt6 where clone E (untreated) and G (resistant to platinum) are sister clones, comparison of E with G and comparison of E with H, a distant but also platinum resistant clone, both showed passage decline in differential expression (Figure 4b). Taken together this indicates gradual transcriptional adaptation to repeated platinum exposure, even within resistant clones.

We next asked whether and to what extent treatment withdrawal influenced the clonal drug resistance transcriptional states (Figure 4a,e-g and Supplementary Figure 9c). In treatment holiday intervals, the majority (79% over all samples) of the DE transcription reverts in the direction of untreated, indicating a collapse back to the earlier state. Notable exceptions are later passages of Pt4 where the tumour cell population exhibited a full CNA clonal sweep of clone A (strongly platinum resistant genotype) at the last two passages, which we have previously shown was the fittest under platinum treatment. An extreme example of loss of transcriptional plasticity after clonal fixation was observed for Pt4, where the later passages X6, X7 of platinum exposure resulted in a monoclonal population [18]. In this case, few reversible genes were observed (only 14 genes at X6 and 19 genes at X7, all *cis*, whereas 1600 genes were found at X5, Supplementary Table 7), consistent with a fixed genomic landscape of drug resistance (Figure 4e-g), with minimal transcriptional plasticity. A similar observation was made for clone A of Pt5, where the transcriptional landscape is fixed at the last passage (X10). On the contrary, clonal sweeps of resistant clones were not detected in Pt6 (see also Figure 2c), and similarly a large number of reversible genes were found at all time points (> 695 for Pt6, Figure 4e). This suggests that these genes develop into new states that are different from untreated and platinum treatment. Taken together, the degree of transcriptional adaptation in single CNA clones on initiation or withdrawal of treatment appears inversely related to the relative contribution of genotypic fitness to platinum.

To define the functional pathways implicated in clone specific and clone independent gene expression, we mapped differentially expressed genes from contrasts between the selected clones (Figure 3a) across temporal passaging, for drug treatment and drug withdrawal versus their untreated temporal counterparts (Figure 5, Supplementary Table 8). We used pathway enrichment tests and Hallmark reference gene sets to highlight the most significant pathways that are related to cancer. We found that the number of regulated pathways for treated versus untreated clones increased with serial exposure for all Pt4,5,6 (from 8 at X4 to 24 at X7 for Pt4, from 8 at X6 to 13 at X10 for Pt5 and from 12 at X5 to 19 at X8 for Pt5, see also Figure 5c). More significant pathways (p < 0.05) are activated at later time points, relative to untreated counterparts.

**Figure 5.**
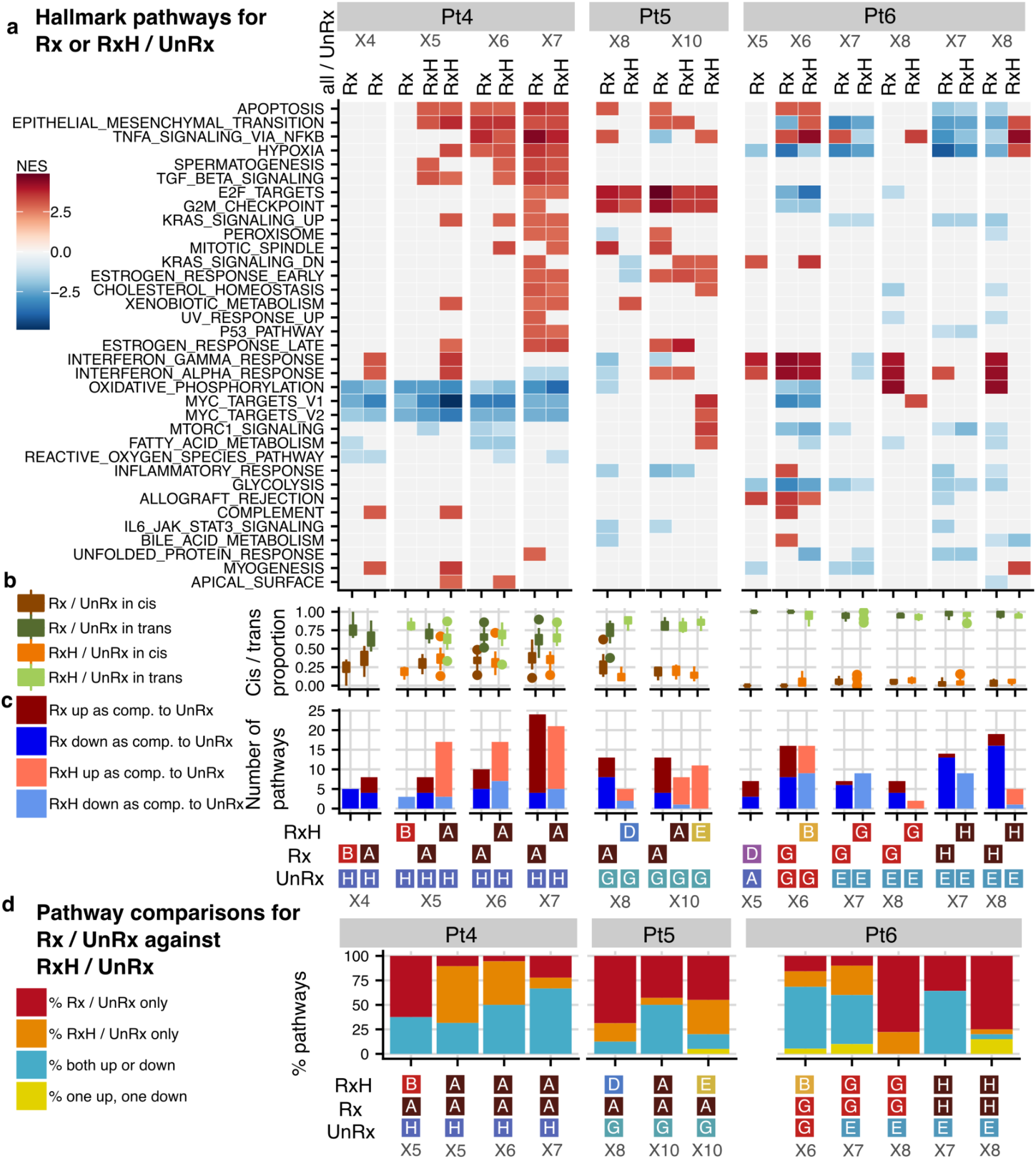
Hallmark Significant Pathways in Rx, or RxH versus UnRx for Pt4-Pt6. (a) Significantly enriched pathways (p < 0.05, vertical axis) from a ranked gene set enrichment analysis (GSEA) [56], using the Hallmark gene set collection from MSigDB [56,57]. Each column corresponds to one comparison between a treated or holiday clone versus an untreated clone (as denoted in Figure 3a), for a specific patient and time point. The color intensity signifies the normalized enrichment score (NES) results of enrichment analysis obtained by using all the edgeR differentially expressed genes for that comparison at FDR < 0.01 and |log2 fold change| >= 0.25. Only the common pathways that are enriched in at least three DE comparisons across all patients are shown. (b) The distribution of cis and trans pathway gene proportions for all the comparisons in (a). (c) The number of up-regulated and down-regulated pathways in each column in (a). (d) Pathway status of treated-untreated comparisons against the pathway status of their respective holiday-untreated comparisons, split into four types of changes: (i) “Rx only” include pathways that are enriched in treated, but not in holiday; (ii) “RxH only” are pathways that were not enriched in treated, but are enriched in holiday; (iii) “both same direction” are pathways that are enriched in the same direction in treated and holiday; and (iv) “both reverse direction” are pathways that are enriched in a different direction in treated and holiday samples.

While Pt4 and Pt5 displayed more up-regulated pathways than down-regulated pathways, especially after several rounds of treatment, Pt6 showed the opposite trend in the last two time points, showing heterogeneity in transcriptional responses across patients. This is consistent with previous work [18], and could be because of different DNA repair deficiency phenotypes of Pt4 and Pt5 as BRCA2 and BRCA1 types respectively, while Pt6 [24] was exhibiting APOBEC-BRCA1 germline mutational signatures. Interestingly, different clones of the same sample may encode common as well as individual pathways, as observed for example in Pt5 X10, where holiday clones A and E display four common pathways and ten individual pathways, when compared against the corresponding untreated sample (Figure 5a).

The cis and trans composition of the genes in the hallmark pathways was consistent with the proportions discussed earlier in Figure 3 (mean percentage of cis genes 34%, 19% and 3% for Pt4,5,6, respectively, Figure 5b). When comparing the enriched pathways for treated versus untreated clones (Rx vs. UnRx) and for drug withdrawal versus untreated clones (RxH vs. UnRx, see the columns in Figure 5a-c), we once again observed the clonal fixation for Pt4, with more similar pathway enrichment at X7 than at X5), but continuous differentiation for Pt6, with the pathway enrichment diverging drastically more at X8 than at X6. Pt5 also displayed no clonal fixation, but less differentiation than Pt6.

We have previously observed that the gene expression regulation may change after withdrawal of drug (Figure 4). Similarly, for our Pt4-6 samples we see pathways that are enriched after drug, but not after drug withdrawal, and vice-versa (Figure 5d, “Rx / UnRx only” and “RxH / UnRx only”, respectively). For treatment induction (“Rx / UnRx only”), a maximum of 60%, 65% and 76% of pathways diverged respectively in Pt4, Pt5 and Pt6. On the contrary, some pathways were enriched only in the holiday versus untreated clones (Figure 5d, “RxH / UnRx only”), with the largest percentage occurring for Pt4 at X5 (52%). The majority of the remaining pathways displayed enrichment in the same direction (both up or both down). For Pt4 and Pt5, the largest percentage of such pathways was observed at the last time point (70% and 50%, respectively), suggesting some degree of pathway fixation over time, while for Pt6, this was observed at the second last time point (70%), suggesting that pathways are still differentiating (Figure 5d). Fewer than 5% of the pathways changed direction of enrichment, from up to down regulated or vice-versa (Figure 5a,d) in any of the contrasts. Taken together, our data suggest that, while some pathways remain stable after drug withdrawal, most either revert to the untreated state, or move into a new state distinct from treated or untreated transcriptomes, indicating an acquisition of transcriptional memory of prior treatment (see trajectory analysis, below).

Finally, we asked which cellular functions are represented by the cis and trans encoded transcription dynamics to drug exposure and withdrawal. Shared common pathways between treated cells (Rx) versus untreated cells (UnRx) across patients Pt4,5,6 include apoptosis, Epithelial Mesenchymal Transition (EMT), Hypoxia, TNF signaling and cell cycle/checkpoint functions (Figure 5). EMT is known to significantly contribute to chemoresistance by converting epithelial cells into mobile mesenchymal cells and altering cell to cell adhesion as well as the cellular extracellular matrix, leading to invasion of tumor cells [25]. TGF-β signaling is known to create pre-target cisplatin resistance and is a critical cellular initiator of EMT [26,27]. Hypoxia is known to create post-target resistance through pro-apoptotic effects and off-target cisplatin resistance [28,29] through high expressions of TMEM45, ID1, ID4 genes (Figure 3c). TNFA signaling via NF-κB develops cisplatin resistance through the activation of mediators, including anti apoptotic genes [30]. This pathway was also found to exhibit reversal of expression on drug withdrawal (Figure 5a). Some pathways exhibited patient specific expression, including MTORC1, IL-2_STAT5 signaling and KRAS signaling down. Pt4 exhibited differential expression of MYC targets including C-MYC, Pt6 exhibited regulation of interferon response pathways implying sensitivity to DNA damage. Notably, Pt6 tumour with an APOBEC mutator phenotype (Pt4, 5 have HRD phenotype), exhibited recruitment and repression of interferon gamma pathways and G2M checkpoints. Furthermore, activation of the interferon alpha signaling pathways could contribute to apoptosis and cellular senescence but are also attributed to increased migration and drug resistance depending on the interferon-stimulated genes transcribed [31,32].

### Pseudotime analysis identifies intermediate differentiation cell states in reversible drug resistance

We next asked whether states of transcriptional resistance induction and reversion could be identified ab-initio. Differential expression analysis tends to summarize a population by a mean value and is therefore less effective in capturing subtle subpopulations that include distinct intermediate differentiation states. To address this we conducted a pseudotime trajectory analysis (Figure 6, Supplementary Figure 10, Supplementary Figure 11) over actual time series samples.

**Figure 6.**
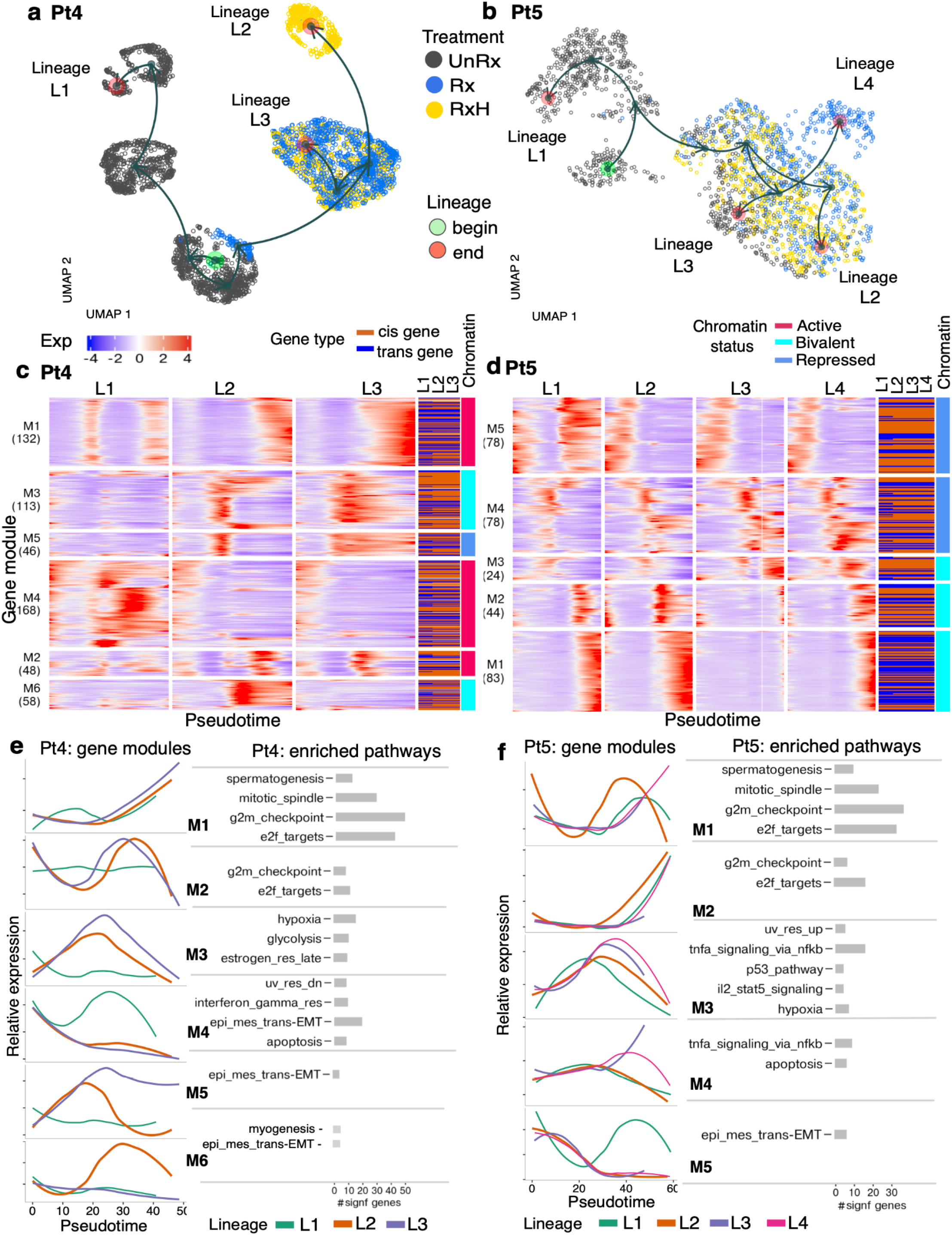
Dynamic gene regulation modules across treatment are captured using pseudotime analysis in Pt4 and Pt5. (a, b) UMAP visualization with individual cell lineages - output of pseudotime analysis for Pt4 (a) and Pt5 (b) coloured by actual drug treatment status (UnRx: untreated cells, Rx: drug treatment, RxH: drug holiday). Green circle denotes starting point of a lineage, red circle denotes the end point of a lineage. (c, d) Heatmap of gene expression across the pseudotime of different lineages for Pt4 (c) and Pt5 (d). X-axis: relative smoothed gene expression of each gene in each lineage, colors denoting gene expression level, blue - low expression, red - high expression. Gene types of each individual gene across different lineages, dark chocolate: cis, blue: trans gene. Chromatin status of each gene module: active - red, bivalent-cyan, repressed-blue. Y-axis: each row is an individual gene. Rows are grouped by regulatory gene modules (M denotes a gene module), i.e. M1(132) means gene module 1 and contains 132 genes. (e, f) Summary of relative gene expression for individual lineages in each gene module from heatmap panels c,d and list of Hallmark enriched pathways related to each gene module based on enrichment analysis gprofilers statistical tests with P-adj<0.05. The bar size denotes the number of genes in each gene module that belong to significant Hallmark gene sets (M6 in panel e did not have any significant gene sets, therefore we included the pathways that had the most number of genes). X-axis: pseudotime, Y-axis: relative expression scaled from -2 to 2.

Firstly, we applied *Slingshot* [33], an efficient computational method that can robustly infer multiple branching cell lineages and pseudotime from single cell gene expression data. For each patient Pt4-6, we used all untreated, treated, and drug holiday scRNA-seq cells (see Table 1), and applied *Slingshot* to obtain inferred lineages over pseudotime. As a result, each cell can be associated with one or more lineages and be assigned a pseudotime value (Figure 6, Supplementary Figure 10, Supplementary Figure 11, Methods).

In Pt4, the pseudotime inference output embedded in UMAP projection reveals two main distinct lineages that correspond to the untreated cell population (lineage L1), and the treated/holiday cell population, confirming the expected differentiation between the treated versus untreated cells. However the treated/holiday population was further sub-divided into two sub-lineages: L3 associated with continuous drug exposure, and L2 associated with drug holiday (Figure 6a). Sub-lineages L2 and L3 share the same starting clusters at the beginning of treatment where cells have a similar effect to the drug, then at the late time points, with enough drug exposure, cells are divided into resistant cell subpopulations (resistant clone A, fitness coefficient median=1.048, sd=0.019 Supplementary Figure 10a,b) along lineage L3, and drug holiday reversibility cells population L2 (drug holiday clone B, fitness coefficient median=1.023, sd=0.015 Supplementary Figure 10a,b) along lineage L2 of Pt4. This result is consistent with the results at the genomic level from our previous study [18] (Figure 2a,c). The UMAP projection (Figure 6a) further suggests that the drug holiday cells in L2 are distinct from the untreated cells, suggesting that the cells exist in a different state than fully resistant or fully untreated. A similar trend was observed for Pt6 (Supplementary Figure 11), where Pt6 lineage L2 appears to represents a distinct treatment holiday state, distinct from the variation between untreated and treated states. As with Pt4, some holiday cells fall back towards the untreated, or remain close to the treated state. In contrast, Pt5 exhibited no distinct holiday state. Here two main lineages that correspond to untreated cells (L1) and treatment/holiday cells, further differentiated into three treatment sub-lineages that are differentiated by cells at the late drug time points (L2, L3) and L4 containing mostly one round of treated cells. Importantly, the cell landscape at the end points of lineage L2 contains drug resistant cells (resistant clone A, fitness coefficient median=1.047, sd=0.018, Supplementary Figure 10c,d). Moreover, the end points of lineage L3 contains drug holiday cells in the similar landscape as untreated cells (Supplementary Figure 10c,d), suggesting that the majority of drug holiday cells revert back to states that are close to the ground state of untreated cells.

We next asked which biological functions are represented in the different lineage states of treatment and treatment withdrawal and their relationship to known states of drug resistance. We used *tradeSeq* [34] to perform between-lineage statistical tests in order to extract differentially expressed genes across lineages in Pt4-6. Moreover, *tradeSeq* admits identification of differentially expressed genes between the end points of each pair of lineages where there was a strong drug effect (see Methods).

The identified *tradeSeq* significant genes (Wald statistics tests assessing differentially expressed genes along multiple lineages, and between early, or end points of lineages, p-value<0.05, see Methods) were grouped into gene modules (Figure 6c,d, Supplementary Figure 11, Supplementary Table 9). The computed gene modules, identified three main patterns of gene regulation with drug effects: (i) genes that are induced relative to untreated state and travel similarly with treatment and holiday (Pt4 modules M1, M2 31.9%; Pt5 M2, 14.3%; Pt6 M4 19.8%); (ii) genes repressed relative to the untreated state by treatment (Pt4 module M4, 29.7%; Pt5 M3, M5 33.2%; Pt6 M2 8.6%) and (iii) strikingly, holiday state genes whose expression is in neither treated nor untreated state, indicating that treatment withdrawal can be associated with novel intermediate states and not simply a reversion to untreated ground state or a fixation to the treated state (Pt4 modules M5, M6 18.4%; Pt6 M3, M5 58.1%). Precisely, genes in modules Pt4 M3, M5 (28.1%) are induced along drug treatment, but are repressed at drug holiday. In contrast, genes in module Pt4 M6 (10.3%) are repressed along drug treatment, but up-regulated at drug holiday time points (examples, Supplementary Figure 10). The dominant biological processes [35,36] in the distinct holiday states was EMT related functions for Pt4 (M5, M6) and a mixture of interferon response and EMT for Pt6 (Figure 6c,d, Supplementary Figure 11, Supplementary Table 9, M3, M5). In contrast, common functions between shared treatment and holiday states were dominated by cell cycle functions such as G2/M checkpoint, mitotic spindle, E2F targets, consistent with the effect of DNA damaging agents in dividing cells. These results match the pathway analysis results in the previous section (Figure 5). With two separate analyses, we see consistent pathway mechanisms that are activated under drug treatment and drug holiday conditions.

Further, we investigated the potential promoter chromatin states of the identified modules in Pt4-6 using the ChIP-seq data for H3K27me3 and H3K4me3 from untreated breast cancer cell line MDA-MB-468 [37]. For Pt4, we found gene modules M1, M2 and M4 to have active promoter chromatin status with significant enrichment of H3K4me3 (P<0.05, ANOVA) and significant depletion of H3K27me3 (P<0.05, ANOVA) compared to repressed (low expressed) genes. While two modules, M3 and M6, had bivalent status having H3K4me3 enrichment (P<0.05, ANOVA), but having similar high levels of H3K27me3 as found at repressed genes (P>0.05, ANOVA test); and lastly M5 gene module was found to have repressed state with the same chromatin state for H3K27me3 and H3K4me3 as found at repressed genes (P>0.05, ANOVA) (Figure 6c, right column, see Methods). In contrast with Pt4, none of the five modules in Pt5 and Pt6 were found to have active promoter chromatin status, but instead had either bivalent/repressed status in Pt5 (Figure 6d, right column, see Methods) or only bivalent status in Pt6 (Supplementary Figure 11). Unique holiday module genes were associated only with bivalent or repressed marks across Pt4,6.

## Discussion

Copy number mutations alter gene dosage potential over hundreds of genes in breast cancer, thus single mutational events have the potential to impact transcription of many genes [10,11]. Here we show how the relationship between gene dosage in subclones of breast cancer and non-genomic transcriptional plasticity impacts transcription in the setting of platinum treatment and platinum withdrawal in multi-clonal triple negative breast cancers. We contrasted three TNBC patient tumours characterized by long interval, multi-year serial passaging [18] and different genomic instability backgrounds, where serial tumour sampling of platinum treatment and platinum withdrawal were available. To separate genotypic and non-genotypic transcriptional dynamics we generated scRNA-seq from serially passaged tumours, assigning cells to clones with a probabilistic model informed by previous single cell genome sequencing determination of CNA clone genotypes. As anticipated, the proportion of differentially expressed transcripts potentially impacted by gene dosage is a reflection of the fraction of genome altered by CNAs and ranges from 5-50%. Whole genome duplication represented only a minority of tumour cells in Pt4-6 (mean 2.86%, sd 1.92%), thus most of the cis variation arises from chromosomal level CNA.

A striking result is that different clones even within a tumour may exhibit very distinct modes of transcriptional landscape fixation. The dynamics of gene expression on exposure to platinum results in a majority increase in transcripts, dominated by non-copy number associated regions of the genome (*trans*). *Cis* region transcripts were correlated with gene dosage. The pattern of transcription on withdrawal appears to partially reflect the degree of clonal selection and fixation in tumour cell population, consistent with previous studies where it has been shown that phenotypic changes are not necessarily associated with genomic changes, but rather transcriptional plasticity plays a major role [16]. The most extreme example in our data is Pt4, which exhibits very little transcriptional variation upon withdrawal platinum by later passages when the entire population is composed of a single drug resistant clone. Pt5 shows an intermediate effect in clonal selection and at least one clone that exemplifies clonal fixation, whereas Pt6 that has the least genome altered, did not exhibit clonally fixed states of transcription. These results emphasize the internal heterogeneity of clones with respect to the contributions of genomic and non-genomic transcriptional plasticity.

A further question relates to the basis of memory in tumour tissues exposed to drugs. Here we observed that while the majority of drug induced transcription can revert, reversion is not always associated with return to the untreated state but rather new states that represent a potential reservoir of intrinsically resistant cells. Perhaps this plasticity results in priming or keeping the memory of exposure [38] through formation of new states that could define the fate of those cells. Pathway analysis shows that convergence on altered EMT state, a known phenotypic background for drug resistance, is one common factor in the reversion trajectories, additionally common interferon pathways and cytokine signaling represent a convergence point.

It should be noted that PDX models, while capturing many aspects of polyclonal tumour evolution and intrinsic tumour cell phenotypes, do not mirror the effects of the human immune system nor replicate precisely the tissue level pharmacology of drug exposure. However the limitations of human tissue biopsy for patients under treatment also preclude the serial analysis needed to measure clonal fitness accurately. The patient tumours studied here represent only a subset of TNBC tumour subtypes, albeit sampled intensively over multi-year intervals. It is likely that recent discoveries of chromosomal instability subtypes of TNBC characterized by foldback inversion patterns (FBI [39]), distinct from homologous recombination deficient TNBC, imply additional mechanisms of cis and trans drug resistance. FBI tumours are intrinsically platinum resistant and tend to exhibit higher genome duplication rates. Nevertheless, taken together our results emphasize that both CNA associated and CNA independent mechanisms may encode transcriptional states associated with platinum resistance and emphasize the internal heterogeneity within tumours with respect to each mechanism. The contribution of each mechanism may have implications for therapeutic strategies. Strong genomic clonal fixation with a clonal sweep implies irreversibility, whereas tumours with heterogeneous clones may amplify non-genomic transcriptional states, with implications for re-treatment or re-sensitization approaches. Future studies of transcriptional plasticity should account for both aspects to achieve a realistic understanding of polyclonal tumour responses to therapy.

## Methods

### Establishment and serial passaging of patient derived xenografts

#### Ethics

The Ethics Committee at the University of British Columbia approved all the experiments using human resources. Patients in Vancouver, British Columbia, were recruited and samples were collected under the tumour tissue repository (TTR-H06-00289). Samples were transplanted to mice under the Animal Resource Centre (ARC) bioethics protocol (A19-0298-A001) approved by the Animal Care Committee (University of British Columbia BC Cancer Research Ethics Board H20-00170) protocols.

Xenograft-bearing mice were euthanized when the size of the tumours approached 1000 mm^3^ in volume (combining together the sizes of individual tumours when more than one was present). The tumour material was excised aseptically, then processed as previously described [18,40]. Briefly, the tumour was harvested and minced finely with scalpels, then mechanically disaggregated for one minute using the Stomacher 80 Biomaster (Seward Limited, Worthing, UK) in 1-2 mL cold DMEM-F12 medium. Aliquots from the resulting suspension of cells and fragments were used for xenotransplants in the next generation of mice.

#### Histopathology of PDX tumours

The hormone receptor status of all tumour samples were determined by immunohistochemistry. Two separate tissue microarrays (TMAs) were prepared using duplicate 1 mm cores extracted from formalin-fixed paraffin-embedded (FFPE) blocks, containing materials from patient derived xenografts (Pt4, Pt5, Pt6) as described in [18,24] (Supplementary Figure 1).

#### TNBC PDX timeseries treatment with cisplatin

Triple negative breast cancer patient derived xenografts were serially passaged as previously described [18,40]. Serially transplanted material represented approximately 0.1 - 0.3% of the original tumour volume. Immunodeficient NOD/Rag1−/−Il2rγ −/− (NRG) mice of the same age and genotype as above were used for transplantation treatment experiments. Drug treatment with cisplatin (Accord DIN: 02355183) was started when the tumour size reached approximately 300 mm^3^ to 400 mm^3^. Cisplatin was administered intraperitoneally (IP) at 2 mg kg^−1^ every third day for 8 doses maximum (Q3Dx8). Low dose cisplatin pulse was selected to achieve the experimental aims of tumour resistance at the time of tumour collection. Residual tumours after treatment were re-transplanted in a new cohort of eight mice. Also, in parallel for the treatment/treatment holiday study group, half of the mice were treated with cisplatin when tumours exhibited approximately 50% shrinkage, the residual tumour was then harvested and re-transplanted for the next passage in the group of eight mice. Again, half of the mice at the second cycle of treatment were kept untreated while the other half were exposed again to cisplatin following the same dosing strategy. Four cycles of cisplatin treatment were generated, with a parallel drug holiday group at each passage.

We used the label Rx to denote treated samples. RxH denotes samples treated over one or more passages but kept on drug holiday (drug withdrawal) at that specific (last) passage. UnRx denotes samples that were never exposed to drug treatment. For patient naming, Pt1, Pt2, Pt3 in this study correspond to the breast cancer patient-derived xenograft SA501, SA530, SA604, and Pt4, Pt5, Pt6 in this study correspond to SA609, SA535, and SA1035 respectively in our previous study [18], see also Table 1 and Supplementary Table 1.

#### Single cell whole genome sequencing and library construction with DLP+

Single cell whole genome sequencing and library construction with (Direct Library Preparation) DLP+ were done as described in [18,41]. Briefly, single-cell suspensions from the tumors were prepared by enzymatic digestion with collagenase/hyaluronidase (Stem Cell Technologies, 07912) enzyme mix in serum-free Dulbecco’s Modified Eagle’s Medium (DMEM) at 37°C with intermittent gentle trituration with a wide bore pipette tip. Cells were stained with LIVE/DEAD Fixable Red Dead Cell Stains (ThermoFisher) and using a cellenONE (Cellenion), single cells dispensed into each well on a nanowell chip containing two unique dual indices. DLP+ sequence analysis, copy number determination and quality control filtering were done as in [18].

#### Processing of patient derived xenografts for single cell RNA sequencing

Viably frozen vial of the PDX tumours was thawed and after washing out the freezing media, the tumour clumps and fragments were incubated with digestion enzymes as previously described [18,41]. The cells were resuspended in 0.04% bovine fetal serum (BSA in PBS). Dead cells were removed using the Miltenyi MACS Dead Cell Removal kit and cells were processed as described in [42]. For library construction, the samples at the same time point were sequenced on the same chips to avoid processing artifacts.

### Data filtering and analysis

#### Phylogenetic tree inference, clone determination and clonal abundance measurements in DLP+

We applied the DLP+ single cell analysis pipeline that was developed in the previous study [18,41] for single cell DNA sequencing datasets. Firstly, the read data from DLP+ single cell sequencing was preprocessed using HMMcopy package version 1.32.0 to provide a copy number profile for each cell with 500KB genomic bin regions [43]. Secondly, we reconstructed phylogenetic trees by applying sitka [19], an efficient Bayesian tree inference method to copy number data. Cells are placed at the terminal leaf nodes in the phylogenetic tree, and cells with high similarity in copy number profiles are placed in the neighbourhood area of the same branch in the tree. Thirdly, based on the output of the phylogenetic tree, clonal populations are identified. The connected components of cells in the neighbourhood area on the phylogenetic tree were identified and cells were divided into clones based on the degree of homogeneity in copy number profiles. Fourthly, the fitness coefficient which denotes a growth potential rate of a given clone along the drug evolution was previously quantified using the fitClone tool [2] for Pt4-6 and used as such in this study.

To simplify the phylogenetic tree for visualization in this study, sitka trees were collapsed and cells in the same clone were represented by one supercell - one circle with the given clone label. The collapsed tree allows us to observe the order of clones, and the clades in the tree. The output of the collapsed trees are shown in Figure 2a. The clones with the fittest coefficients across treatment as calculated by fitClone are annotated with *Rx, and across untreated lines are annotated with *UnRx. The phylogenetic trees of the three drug treated patients Pt4, Pt5, Pt6 were generated in our previous study [18]. We applied the same methods to reconstruct the trees for three untreated patients Pt1, Pt2, Pt3 in this manuscript (Figure 2a).

#### Quality control for the scRNA-seq data

Firstly, count matrices were generated using CellRanger version 3.0.2 (V3 chemistry), see column “Counts_sequenced” in Supplementary Table 1. Secondly, all mouse cells that were mixed with human cells were eliminated. Cells were aligned using both mouse and human reference gene sets. A cell is classified as a mouse cell if the total number of counts aligned to mouse reference of the 10x sample was greater than the total number of counts aligned to the human reference (column “Counts_no_mouse_cells”). Thirdly, cells were considered to have passed a quality control filter (QC-filter) and retained for subsequent analysis if they met the following criterion: (i) minimum of 1000 detected genes, (ii) low mitochondrial contamination with less than 20% of UMIs counts mapping to mitochondrial genes, (iii) less than 60% of UMI counts mapping to ribosomal genes, and (iv) the total counts (UMIs) per cell was at most 3 median absolute deviations lower than the overall median counts. Cells with lower quality than the above criteria were filtered using the calculateQCMetrics and isOutlier functions in the scater package [44] (column “Counts_quality_cells”). Fourthly, we eliminated doublets using package scrublet [45] (column “Counts_no_doublets_final” in Supplementary Table 1).

#### Clonealign

Clonealign version 1.99.2 [20] was used to align scRNA-seq cells from each specific xenograft sample to the DLP+ clones obtained from the same xenograft sample. Clonealign assumes that, for most genes, the non-diploid copy number is positively correlated with the gene expression level. Manual tuning was necessary for some of the parameters due to the high degree of heterogeneity in the noise of our samples (see Supplementary Table 3), as follows:

1. Copy number purity threshold for a gene in a clone, defined as the percentage of cells that have the modal copy number for that gene and clone, was set to 0.6 for most samples. For example, if the copy number mode of a gene in a clone is 4, then the gene is included in the clonealign model only if at least 60% of the cells in that clone have copy number 4 for that gene. This ensures that only the highest confidence data is included in the analysis. However, for some samples from Pt4 and Pt6, no or very few genes pass this threshold, therefore we used lower thresholds for those samples.
2. initial shrink, defined as the strength with which the variational parameters for clone assignments are initially shrunk towards the most likely assignments, was set to 0 or 10.
3. data init mu was set to TRUE if the mu parameters need to be initialized using the data, or FALSE otherwise.

Other parameters that had the same values for all libraries include: n repeats = 3, mc samples = 1, learning rate = 0.07, max iter = 500, saturation threshold = 6 and clone call probability = 0.9 (a cell is assigned to a clone if the posterior probability is at least 0.9, otherwise it is unassigned).

#### Differential expression analysis

Differential expression quantifiers including log2 fold change and false discovery rate (FDR) were computed using the R 3.6.0 Bioconductor package edgeR 3.26.0 [21] that implements scRNA-seq differential expression analysis methodology based on the Negative Binomial distribution. First, we applied scran normalization to remove library size effects and use the obtained size factors for the next step. Then, we called the *estimateDisp* function to estimate the dispersion by fitting a generalized linear model that accounts for all systematic sources of variation. Next, we used the edgeR functions *glmQLFit* and *glmQLFTest* to perform a quasi-likelihood dispersion estimation and hypothesis testing that assigns FDR values to each gene. In the track scRNAseq plots in Figure 3 and Supplementary Figure 5, a positive log 2 fold change value for Clone X relative to Clone Y signifies that the gene is significantly more up-regulated (at a given FDR threshold) in X than in Y while taking into consideration all the expression values for all the genes in both clones. Similarly, a gene with negative log 2 fold change is significantly more down-regulated in X than in Y.

#### Normalization method

We applied SCTransform [46] to normalize the gene expression matrix and regress the difference in sequencing depths between samples. The normalized matrix will be used for pseudotime analysis and UMAP visualization. First, scRNA-seq data were preprocessed. Low expression genes that have zero UMI counts in more than 97.5% of the cells were considered as unreliable genes and were excluded from our analysis. Then, from the list of filtered genes, the mitochondrial confounding genes were removed. SCTransform model genes using Pearson residuals from regularized negative binomial regression. The generalized linear model from SCTransform uses a covariate value that accounts for sequencing depth. First we removed genes with low expression and confounding genes as described above. Then we used the SCTransform function within Seurat version 4.0 to normalize data, resulting in a SCTransform log-normalized expression matrix.

#### Dimensionality reduction computation

To visualize the dimensionality reduction map, we applied a basic processing pipeline in Seurat version 4.0 [47] to compute the UMAP features map. First, the gene expression matrix was normalized as described above using SCTransform. The 30 principal components vectors (PCA) are computed from a log normalized gene expression matrix. Then UMAP vectors were computed based on 30 PCA vectors using the uwot package that is included within the Seurat version 4.0 platform.

#### CNA genotype dependent in cis, and independent in trans gene detection

To classify genes based on their copy number (CNA) genotype dependent/independent, we implement the following steps: (i) summarize the copy number profile of each clone; (ii) assign copy number profile to genes in each clone at the overlapping of gene genomic regions and bin genomic regions; (iii) classify genes into different gene types based on copy number alterations.

i. Computing the median copy number profile of each clone: first we grouped cells based on cell clonal labels achieved from DLP+ analysis procedure above, then for each clone, copy number values at each bin genomic region were summarized by the median copy number value of all cells at this bin genomic region in the given clone.
ii. Assigning copy number profile to ensembl gene indexes: to find mapping between DLP+ copy number results and scRNA-seq gene expression data, all overlaps between bin genomic regions and the genomic ranges of transcriptomic genes were taken into account. We used the R package org.Hs.eg.db version 3.8.2 [48] to find all genomic ranges (containing start, end, width) corresponding for a given ensembl gene index. Then findOverlaps function from R package iRangers [49] to find any overlaps between bin genomic regions and genomic ranges of each gene, resulting in the list of bin genomic regions and their corresponding gene indexes at the overlapping area.
iii. Classifying genes into CNA genotype dependent/independent cis, trans gene types based on copy number alterations: the input data for gene classification is differentially expressed genes in scRNA-seq data between two clones that were inferred using clonealign and copy number alteration between two clones from DLP+ analysis at the overlapping genomic region. CNA genotype independent in cis gene is a gene that exhibits the change in gene expression (DE gene) in scRNA-seq gene expression, and its copy number value. In contrast, CNA genotype independent in trans gene is a gene with its change in expression that only occurs at the transcriptomic level but does not exhibit any change in copy number values at the overlapping genomic regions. To compute cis genes, from DE genes that were obtained using edgeR differential expression analysis as described above, we tracked the assigned copy number profile for each gene, in case there is a change in copy number values, these genes were labelled as in cis genes, otherwise were labelled as in trans genes.

#### Gene classification: define six variation trends

Four main variation trends for cis genes based on the positive or negative directions of gene expressions and copy number profiles, and 2 variation trends for trans genes based on the direction of gene expressions were defined. Four main variations in cis genes are: (i) Gain Up: gain, increase in median copy number values between two clones in DLP+ results and up-regulated with positive logFC value of gene expression in DE analysis between two inferred clones in scRNA-seq. (ii) Loss Down: loss, decrease in copy number values between two clones in DLP+ results and down-regulated in gene expression with negative logFC value of gene expression in DE analysis between two inferred clones in scRNA-seq. (iii) Gain Down: gain, increase in median copy number values and down-regulated in gene expression (iv) Loss Up: decrease in copy number values and up-regulated in gene expression. Moreover, based on the directions, we classified in cis genes into the copy-number correlated in-cis gene in case the gene shows the same direction in both transcript and genomic levels: Gain Up, Loss Down, or copy-number anti-correlated in cis genes in case opposite direction: Gain Down, Loss Up. Two variation trends that can occur in trans genes are (i) Trans Up: up-regulated gene in scRNA-seq gene expression (U) and (ii) Trans Down: down-regulated gene in scRNA-seq gene expression(D) (Fig3).

#### Mapping reference genes to our DE genes results

Next, we investigated whether the significant genes obtained from our analysis belong to the list of cancer functional genes.To address this, first, we scanned two databases and denoted as custom reference genes sets, including ADAM PanCancer core cancer fitness [23], and the genes involved in cisplatin resistance were curated from recent literature (Supplementary Table 6). Then we computed the fraction of cis, trans DE genes from our results that belong to two reference gene sets (Figure 3, Supplementary Figure 6). Moreover, to examine whether the differentially expressed genes obtained from our analysis enriched any custom reference gene set of core cancer fitness or cisplatin related genes, we adopted the pathway enrichment analysis applying R package fgsea version 1.22.0 [50]. Similar to pathway analysis using well-known gene sets, here we used two custom gene sets as input, and logFC values of significant genes from DE analysis results as input. The significant pathway results with p-adjusted value<0.05 are shown in Figure 3d.

#### Track plots

Top plot used scatter plot in R package ggplot2 version 3.3.3 [51], each dot denote a DE gene, yaxis: log2 fold change value of each gene in DE analysis result, xaxis: the genomic position of gene, calculated by extracting start, end, and chromosome genomic position of a given gene using R package annotables version 0.1.91 (https://github.com/stephenturner/annotables). We divide genes into sub plots based on the chromosome positions (chr 1:22 and chr X) Bottom: heatmap plots of median copy number profile for pair of clones from output of DLP+ copy number analysis.

#### Gene dynamic level quantification

Two types of dynamic genes were identified as follows. Firstly, to identify genes that were induced or repressed following cisplatin treatment (Figure 4a-d), we selected the genes that were differentially expressed between Rx and UnRx (FDR <0.01, |log2 fold change| > 0.5) and non-differentially expressed between Rx and RxH (FDR > 0.1), to obtain a set of genes with significant change in expression after treatment, but stable after drug withdrawal. Some of these genes had increased expression after treatment (induced), and some had decreased expression (repressed).

Secondly, to identify the genes that displayed dynamic changes following drug withdrawal, we computed the differentially expressed genes between Rx and RxH (FDR < 0.01, |log2 fold change| > 0.5), and intersected them with all the UnRx genes in order to measure the direction of change. While most genes reverted back towards the untreated state, some genes changed away from the untreated state.

#### Pathways analysis

Pathway Enrichment Networks were computed from differentially expressed genes (FDR < 0.01) ranked by log2 fold change. A normalized enrichment score (NES) was calculated from a ranked gene set enrichment analysis (GSEA) [34] performed on each subset of differentially expressed genes using the hallmark gene set collection from MSigDB [35]. Significantly enriched pathways (adjusted p-value < 0.05) and pathway specific differentially expressed genes were included in the network enrichment Figure 5.

#### Trajectory analysis

Slingshot [33] is an efficient method which can robustly infer cell trajectories and accurately capture pseudotime. In this study, Slingshot was used to infer pseudotime, and tradeseq [34] was used to extract differentially expressed genes across multiple lineages of pseudotime output. To prepare input data, we applied Seurat v4.0 [46] preprocessing steps. First, from SCTransform normalized gene expression matrix as described above, we selected 3000 most highly variable genes (HVG) using FindVariableFeatures function and ‘vst’ method within Seurat. Following the common pseudotime analysis pipeline, we only take into account the top 3000 HVG genes in this study. The lowly expressed genes do not show the strong regulation, and are excluded from analysis. The normalized expression matrix of HVG was used as input to extract 30 principal component (PCA) vectors. Thus, PCA vectors were used to do cell clustering using the Louvain algorithm, dividing cells into multiple clusters. Here we used the resolution 0.3 to get cell clusters at the fine grain level. Similar to the pseudotime analysis pipeline in general, we removed a small outlier cell cluster, and kept the main cell population.

To run slingshot, first one cell cluster was selected as the starting point of trajectories. In this study, the cluster with the most number of untreated cells from the earliest passage was selected as the starting point. Then, in order to estimate the global lineage structure by building a minimum spanning tree, we called the getLineages function from slingshot using as input 30 PCA vectors of all cells in each series. Based on the graph of main lineages, the smooth branching lineages and pseudotime values are estimated by fitting simultaneous principal curves using the getCurves function from Slingshot. The output is the pseudotime values that are assigned to each cell, and with multiple lineages, cells can be assigned to one or several lineages of development.

To visualize pseudotime results, we projected output into 2 UMAP vectors using embedCurves function (Figure 6a,b, Supplementary Figure 9). Each lineage includes several clusters in the order of pseudotime. We added the arrows from one cluster to the next consecutive cluster in the given lineage to show the direction of development. The starting cluster is marked by a green circle and all intermediate clusters are marked by black circles and the ending cluster of each lineage is marked by a red circle.

We characterized each lineage based on the treatment conditions and inferred clonal cell labels from clonealign results. The prevalence plots of annotated labels were shown in the Supplementary Figures 10, 11.

Based on the output of trajectory inference, we applied tradeSeq in order to detect genes that displayed a strong differentiation between lineages. First, we ran tradeSeq fitGAM function to fit a negative binomial generalized additive model (NB-GAM) to the normalized counts gene expression matrix, pseudotime value of each cell, and cell weights. Output of trajectory inference shows the substantial difference between untreated lineage and drug treatment lineages. In this study, we focus on ‘between lineages’ comparison. A gene that is significant along a trajectory should satisfy two conditions: (i) gene expressions are varying along at least two or multiple lineages applying a statistical test patternTest function in tradeSeq; and (ii) gene expression be significantly different between endpoints - diffEndTest or early dividing points - earlyDETest of two lineages. We used a p-adjusted threshold value of 0.05 to select significant genes. Then, from the list of significant genes, we applied a threshold of 200 for patternTest significant level, and a threshold of 50 for diffEndTest, or earlyDETest to retrieve the most significant genes (Figure 6c,d, Supplementary Figure 10, Supplementary Figure 11).

In order to divide significant genes into multiple regulated gene modules, where each module displays a similar gene regulation pattern, we did gene clustering based on the Monocle3 genes clustering method [52]. From the list of significant genes, the cosine distance metric was applied to compute the distance between genes, and then extract 25 UMAP vectors. UMAP vectors were then used as input to the Leiden clustering method, with resolution 0.3 to extract gene module labels (Figure 6c,d, Supplementary Figure 10, Supplementary Figure 11).

Significant genes along trajectories are classified into cis, trans gene types in each lineage based on the list of inferred clonal labels and copy number profiles of each clone in DLP+. We assessed the inferred cell clone labels in each lineage. The copy number profile of these genes across clones are examined to see whether there is any change in copy number values. If there is a change, the gene is classified as cis gene, otherwise as trans gene in a given lineage (Figure 6c,d, Supplementary Figure 10, Supplementary Figure 11).

The regulation pattern of each gene module is summarized using the geom_smooth function in ggplot2 (Figure 6 e,f, Supplementary Figures 11d). To define the significant hallmark gene set that are related to each gene module, we applied gprofilers statistical tests g:GOSt test using default parameter settings and hallmark reference gene set. The output of the gene sets along with a number of significant genes were displayed in Figure 6e,f and Supplementary Figure 11d.

#### Chromatin status analysis

ChIP-seq data in untreated cancer cell line, MDA-MB-468, for H3K27me3 (GSM2258886 & GSM2258887), H3K4me3 (GSM2258892 & GSM2258893) and input control (GSM2258900 & GSM2258901) was retrieved from GEO [37]. ChIP-seq reads were mapped to the human genome (hg38) using Bowtie2 [53]. Using featureCounts (Subread package v 2.0.3) mapped reads were assigned to the promoters (2kb window upstream of TSS) corresponding to the genes identified as part of the expression modules for each TNBC line in the pseudotime analysis. ChIP/input reads per million was calculated for each promoter in the respective modules to compare the enrichment of H3K27me3 and H3K4me3 with respect to a set of 500 repressed genes with non-zero lowest number of reads in Pt4 bulk data across time-series as a control for the genes with repressed chromatin status. Modules with significant enrichment of H3K4me3 (P<0.05, ANOVA) and significant depletion of H3K27me3 (P<0.05, ANOVA) compared to low expressing or repressed genes were labeled as having ‘Active Promoter’ status. While, modules having significant enrichment of H3K4me3 (P<0.05, ANOVA test), but similar high levels of H3K27me3 as found at repressed genes (P>0.05, ANOVA test) were labeled as having ‘Bivalent Promoter’ status. Lastly, modules were labeled to have ‘Repressed Promoter’ status if they have similar levels of H3K4me3 (low) and H3K27me3 (high) compared to repressed genes (P>0.05, ANOVA).

## Acknowledgments

This project was supported by the BC Cancer Foundation at BC Cancer. Samuel Aparicio holds the Nan and Lorraine Robertson Chair in Breast Cancer and is a Canada Research Chair in Molecular Oncology (950–230610 and CRC-2021-00205). Additional funding was provided by a Terry Fox Research Institute grant (1082), a Canadian Cancer Society Research Institute Impact program grant (705617), a CIHR grant (FDN-148429), Breast Cancer Research Foundation awards (BCRF-18-180, BCRF-19-180, BCRF-20-180 and BCRF-21-180), the Cancer Research UK Grand Challenge Program (C31893/A25050) and the Canada Foundation for Innovation (40044) to Samuel Aparicio. We thank Beixi Wang for their work on single cell sequencing wet lab experiments.

