## Supplementary Figures for "Single cell decoding of drug induced transcriptomic reprogramming in triple negative breast cancers"

[6] CRUK Grand Challenge IMAXT Team

[7] Lunenfeld-Tanenbaum Research Institute, University of Toronto, Toronto, ON, Canada

[8] Department of Molecular Genetics, University of Toronto, Toronto, ON, Canada

*All these authors contributed equally

### Corresponding author

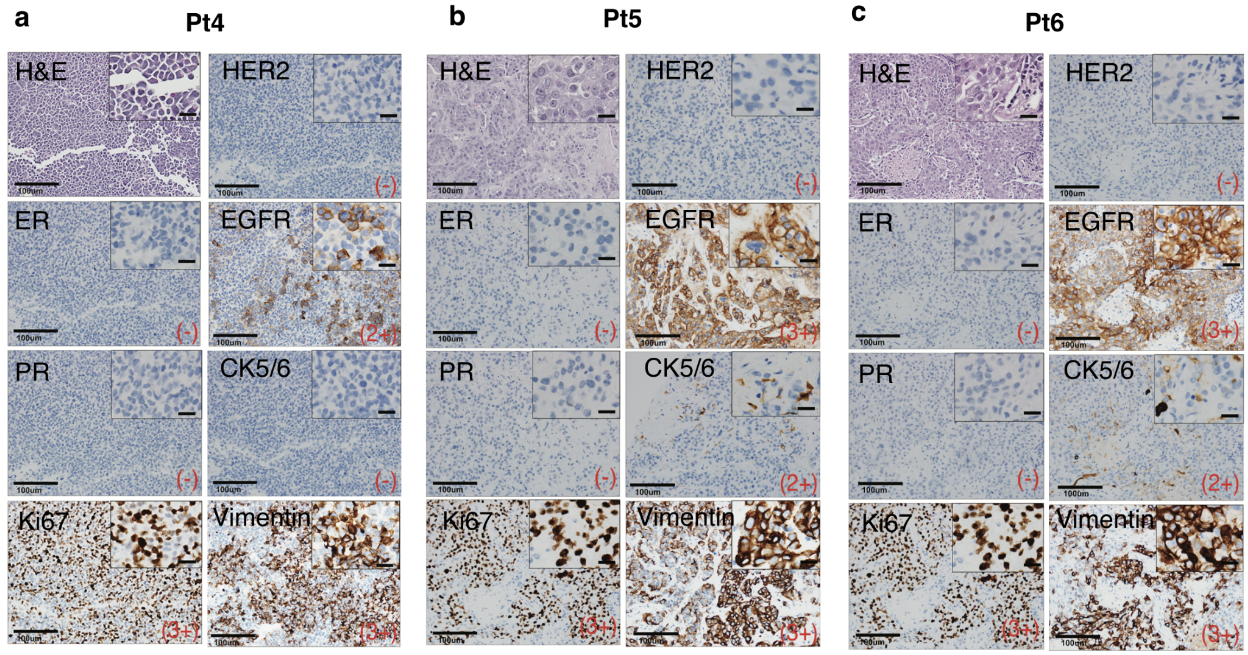


**Supplementary Figure 1: Histology images of TNBC patient's tumor samples with H&E and immunohistochemistry staining** (a) Pt4 (b) Pt5 (c) Pt6. Scale bars 500um (main) and 100um (inset). Right bottom red values, pathologist determined, discretized score ranging from - (no staining) to 3+, based on staining intensity level of stained cells and number of cells that are expressed/ stained a marker compared to total cells in tissue.

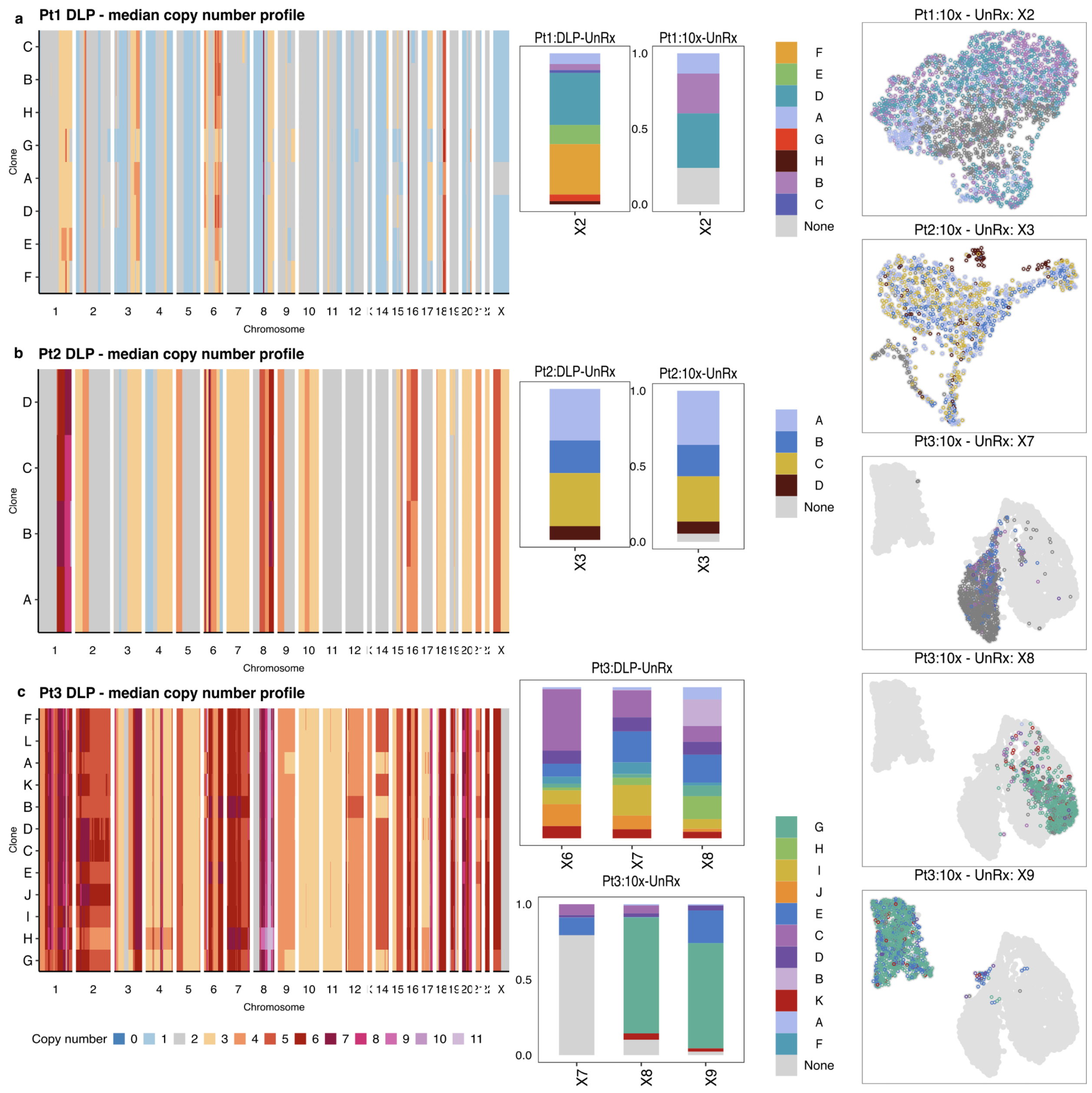


**Supplementary Figure 2: Genomic and transcriptomic characteristics of three untreated patients Pt1-3 (a, b, c).** Segment copy number profiles of Pt1, Pt2, Pt3 in DLP+ sitka phylogenetic tree results at left panels: heatmap with x-axis lists each chromosome, y-axis lists the sitka phylogenetic clones. Each entry represents the median copy number at the corresponding bin genomic position for each clone. Middle panels: Bar plots showing the clonal fractions in DLP+ and 10x data at each time point. Unassigned cells were denoted as None.

Right panels: cell populations UMAP plots in 10x scRNA-seq data, and each dot is one cell, colored by assigned clones.

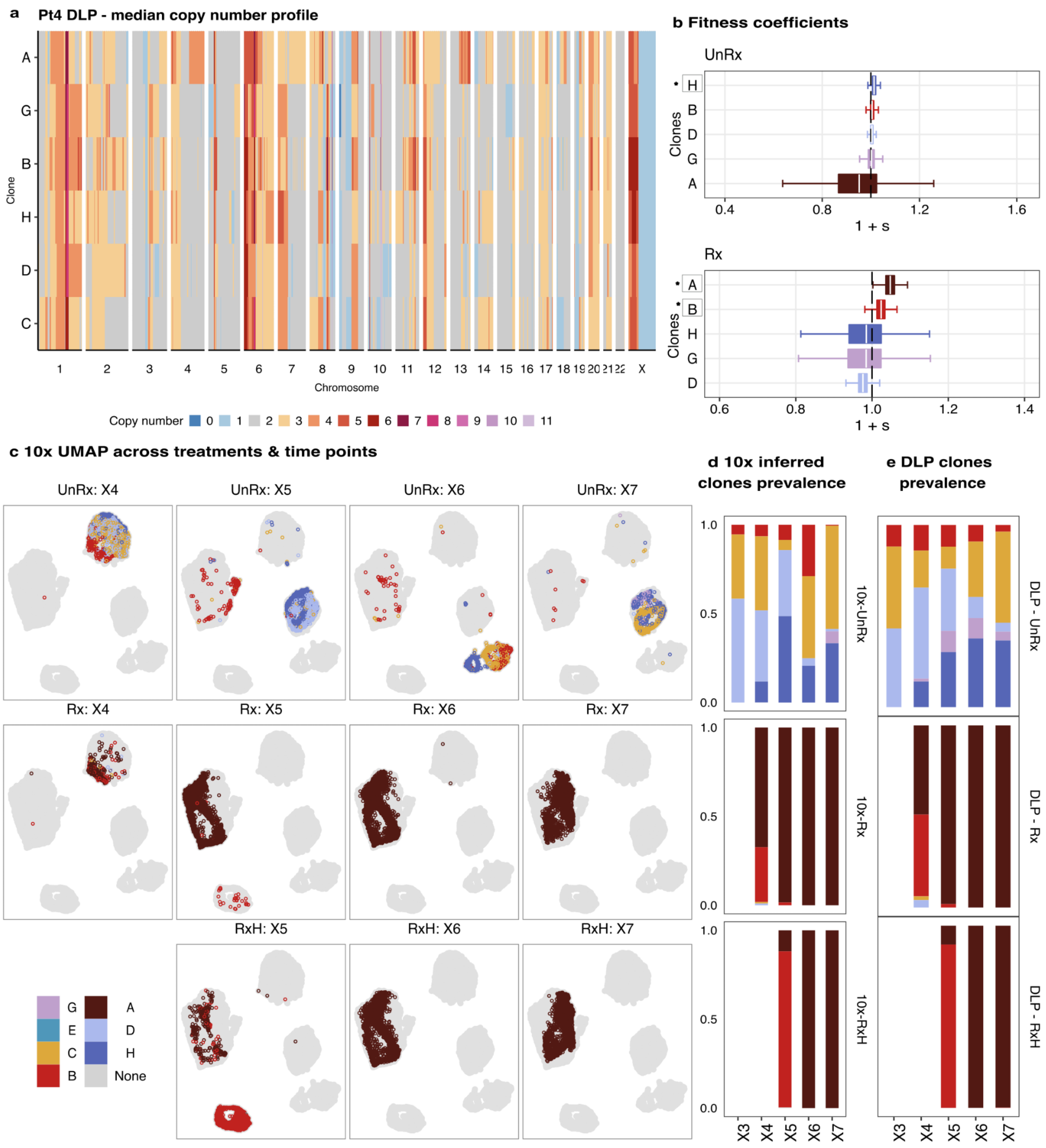

**Supplementary Figure 3**: **Genomic and transcriptomic characteristics of Pt4.** (a) Segment copy number profiles of Pt4 in DLP+ sitka phylogenetic tree results, heatmap with x-axis lists each chromosome regions, y-axis lists the sitka phylogenetic clones. Each entry represents the median copy number at the corresponding bin genomic position for each clone. (b) fitness coefficient of each clone in DLP+ from previous analysis (Salehi et al. 2021). The stars * mark the clones with highest fitness coefficient in untreated, and treated cells. (c) UMAP of 10x scRNA-seq data denote the landscape of cells across drug treatments and untreated time points. Clonal prevalence of inferred clones in scRNAseq analysis (d), and of clones from sitka phylogenetic results in DLP+ analysis (e).

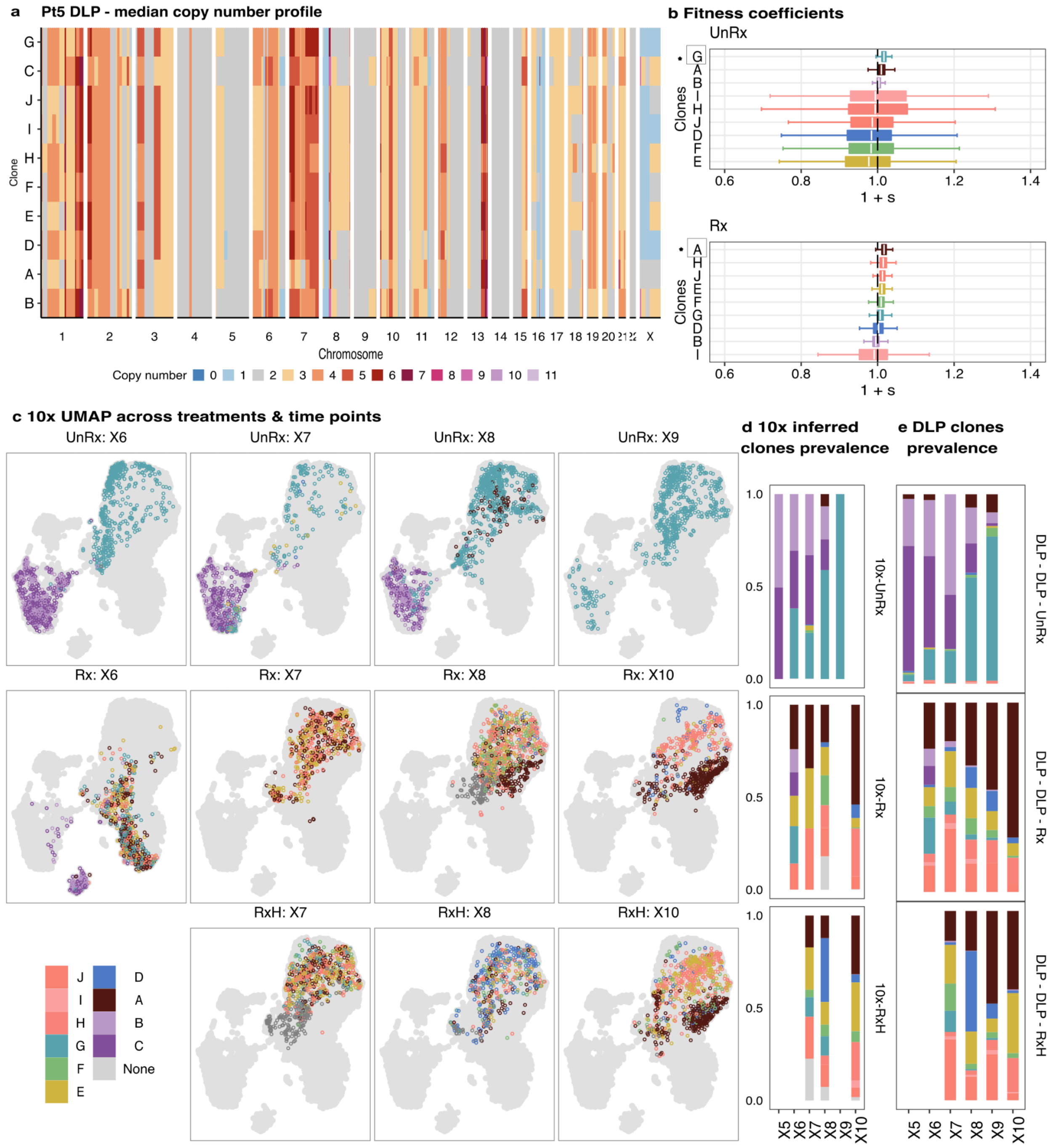


**Supplementary Figure 4**: **Genomic and transcriptomic characteristics of Pt5.** (a) Segment copy number profiles of Pt5 in DLP+ sitka phylogenetic tree results, heatmap with x-axis lists chromosome regions, y-axis lists the sitka phylogenetic clones. Each entry represents the median copy number at the corresponding bin genomic position for each clone. (b) fitness coefficient of each clone at genomic DLP+ data from previous analysis (Salehi et al. 2021). The stars * mark the clones with highest fitness coefficient in untreated, and treated cells. (c) UMAP of 10x sc-RNAseq data denote the landscape of cells across drug treatments, drug holiday and untreated time points. Clone prevalence of inferred clones in scRNA-seq analysis (d), and of clones from sitka phylogenetic results in DLP+ analysis (e).

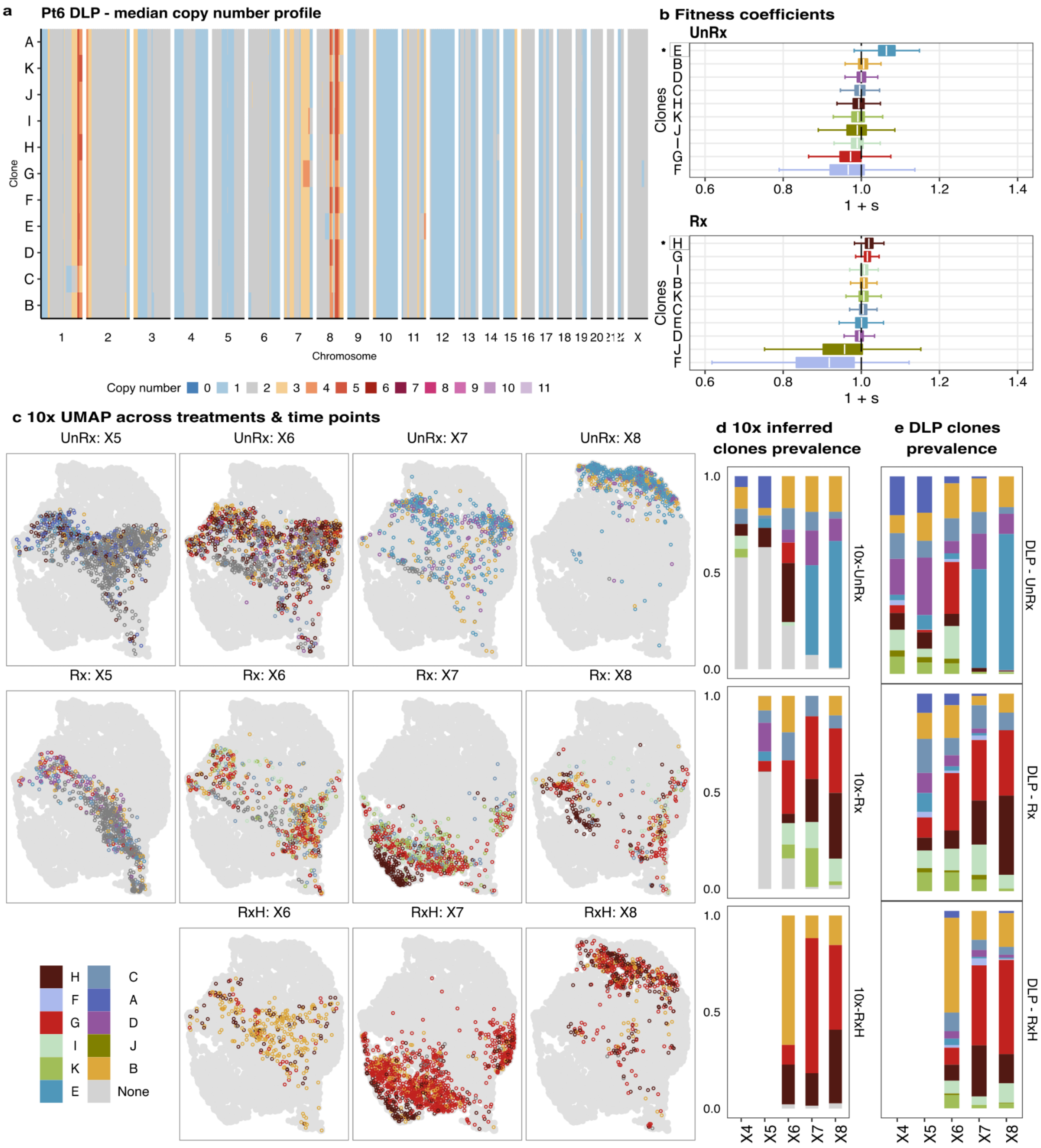

**Supplementary Figure 5**: **Genomic and transcriptomic characteristics of Pt6.** (a) Segment copy number profiles of Pt6 in DLP+ sitka phylogenetic tree results, heatmap with x-axis lists chromosome regions, y-axis lists the sitka phylogenetic clones. Each entry represents the median copy number at the corresponding bin genomic position for each clone. (b) fitness coefficient of each clone in DLP+ from previous analysis (Salehi et al. 2021). The stars * mark the clones with highest fitness coefficient in untreated, and treated cells. (c) UMAP of 10x scRNA-seq data denote the landscape of cells across drug treatments, drug holiday and untreated time points. Clone prevalence of inferred clones in scRNA-seq analysis (d), and of clones from sitka phylogenetic results in DLP+ analysis (e).

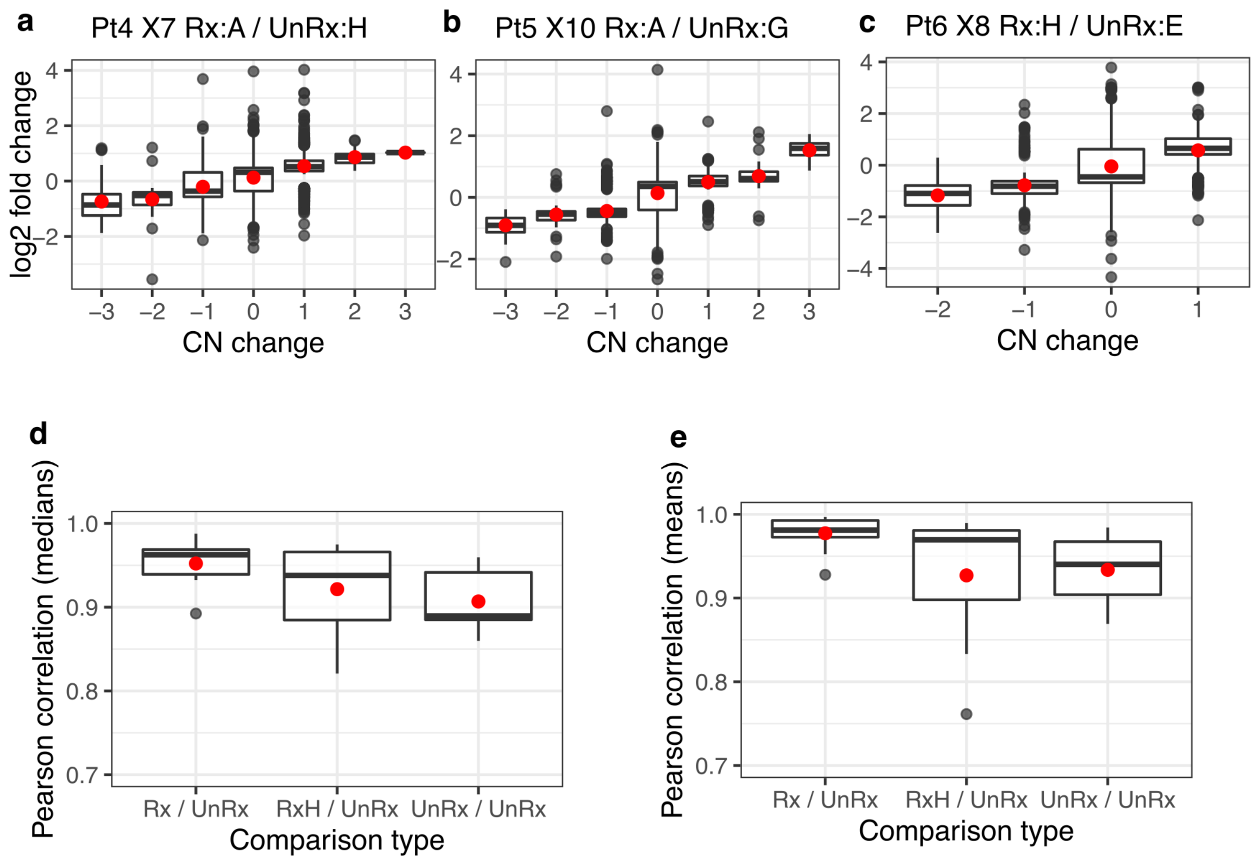


**Supplementary Figure 6**: **Correlation between copy number (CN) change and differential expression.** (a) log2 fold change for the corresponding CN change in Pt4 passage X7 when comparing the differentially expressed genes at Rx:A vs. UnRx:H, as in Figure 3c. The red dots depict means, the black horizontal lines depict medians. Pearson correlation for the medians versus CN change (Pmedians) = 0.99. Pearson correlation for the means versus CN change (Pmeans) = 0.99. (b) Same as a, but for Pt5 passage X10, Rx:A versus UnRx:G, as in Supplementary Figure 7a. Pmedians=0.97, Pmeans=0.98. ( c) Same as a, but for Pt6 passage X8, Rx:H versus UnRx:E, as in Supplementary Figure 7b. Pmedians=0.95, Pmeans=0.99. (d) Boxplots of Pearson correlations for all DE comparisons in Figure 3d and Supplementary Figure 8c,d, by comparison type, using Pmedians. (e) Same as e, but using Pmeans.

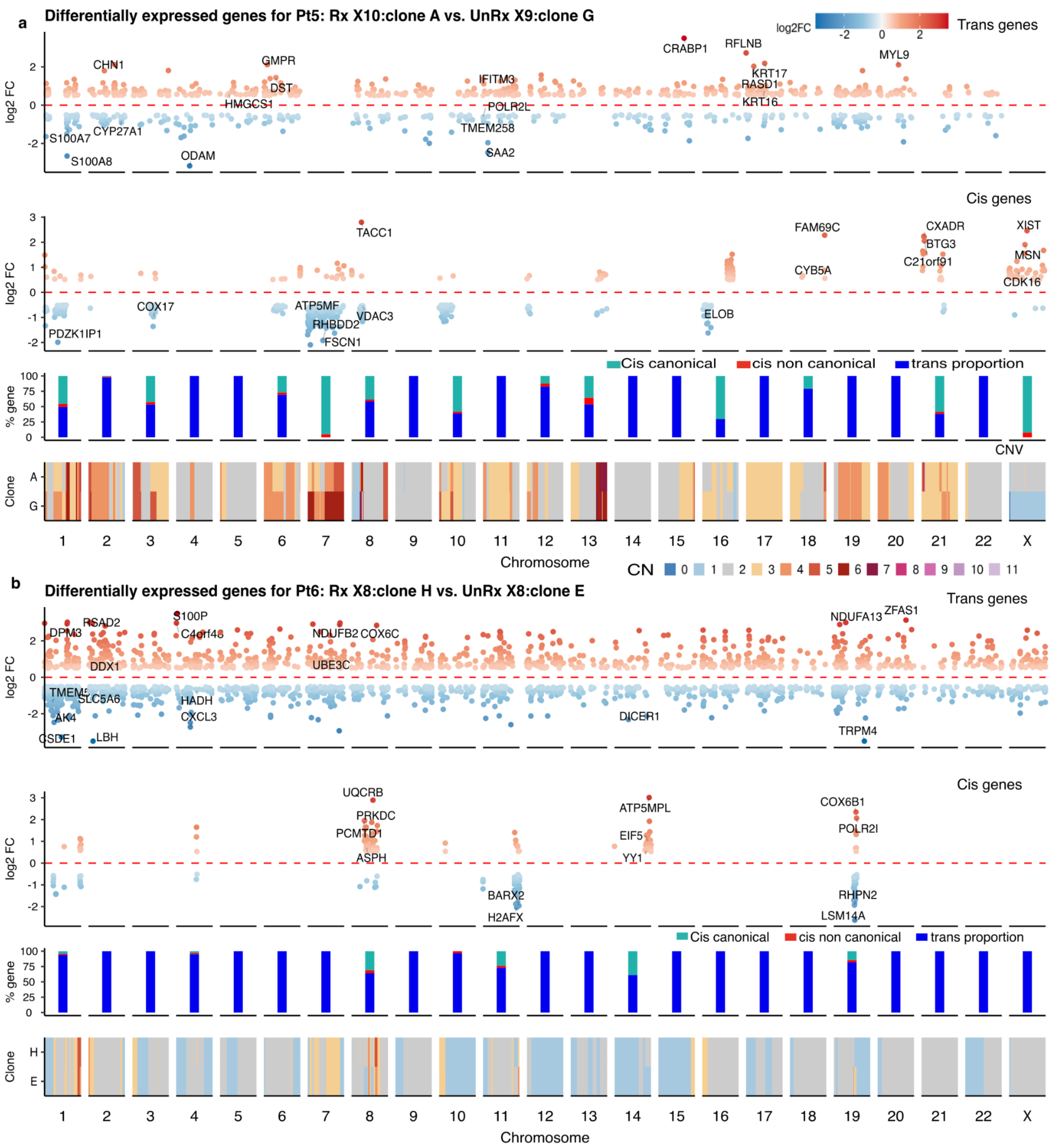


**Supplementary Figure 7**: **Track plots for Pt5 (a) and in Pt6 (b) showing the overlapping regions between gene genomic regions and copy number bin genomic regions**. In each panel, log2 fold change values of trans DE genes (top), cis DE genes are shown. Red and blue gradient colors denote the degree of log2 FC in positive and negative directions, each dot is one DE gene with selected condition abs(log2FC)>0.5, FDR<0.01, p value<0.05. And (third panel)

the distribution of each gene type across chromosomes (green: cis genes canonical, red: cis genes non-canonical, blue: trans genes). At bottom, median copy number of each clone at bin genomic regions across chromosomes.

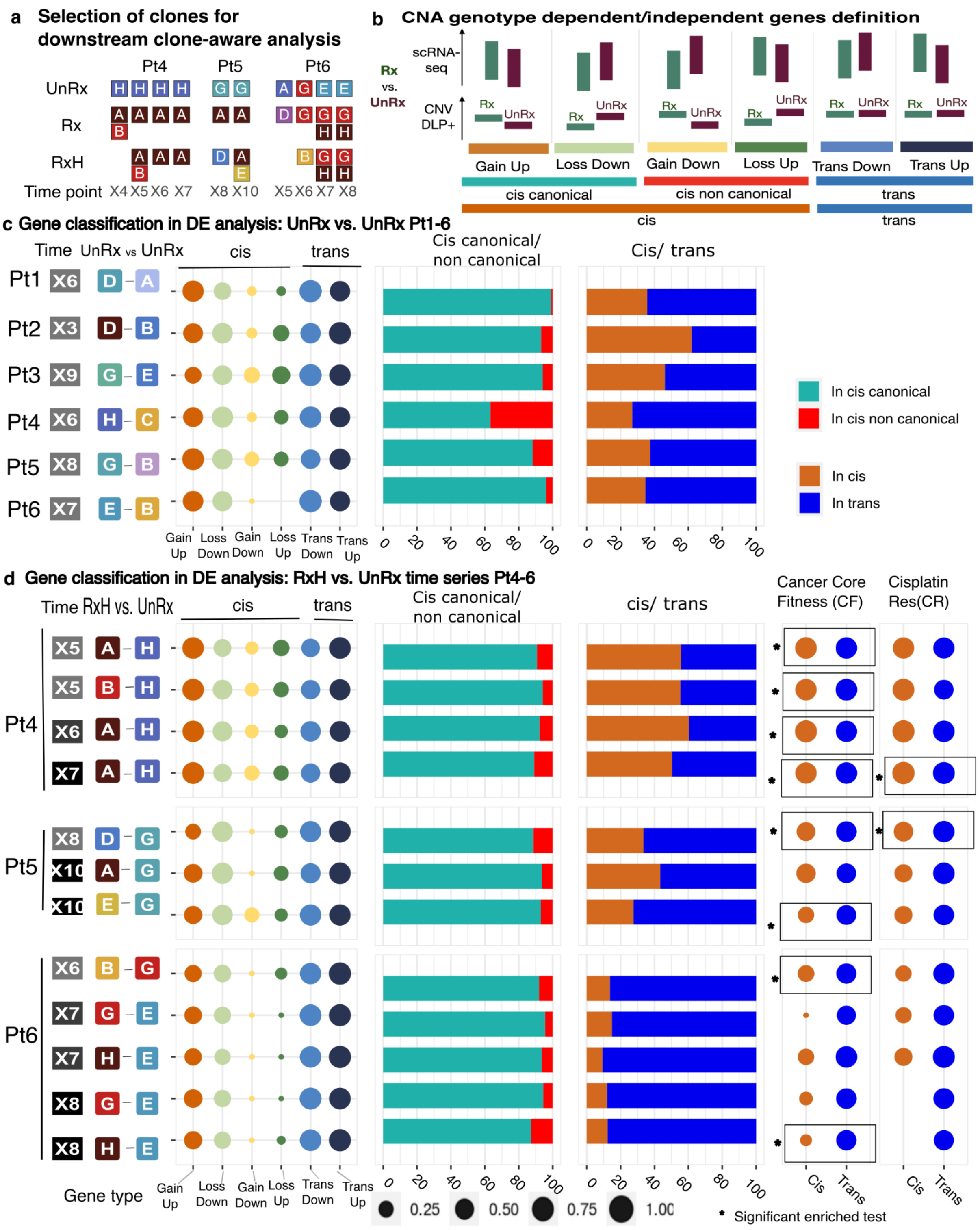


**Supplementary Figure 8: Cis and trans gene proportions and gene set memberships for untreated series, and drug holiday time series.**

(a)For Pt4-6, the clones that were the fittest or most abundant and had at least 100 scRNA-seq cells were selected for further clone-aware analysis.

(b) Gene classification: genes are divided into cis - CNA genotype dependent genes, and trans - CNA genotype independent genes based on whether the copy number values changed at the overlapping genomic regions of the pair clones comparisons. Further, cis genes are divided into canonical and non canonical based on the same or opposite directions of gene expression and copy number values. Genes were then classified based on the direction of change in gene expression (up- or down-regulated) as compared to the direction of change in copy number (gain or loss).

(c) Differentially expressed genes between UnRx versus UnRx for Pt1-6 between the two most dominant clones in untreated patients Pt1-3 and untreated Pt4-Pt6. The dot size represents the proportion of each gene type based on the definition in b: cis canonical (light blue) and cis non canonical (red), cis genes (chocolate) and trans genes (blue).

(d) Differentially expressed genes between drug holiday clones versus sensitive clones (RxH versus UnRx) in Pt4-6. The dot size represents the fraction of each gene type based on the definition in (b). Cis canonical (light blue) and Cis non canonical (red), cis genes (chocolate) and trans genes (blue). Gene set membership: mapping of cis and trans genes to two reference gene sets: COSMIC core fitness census genes (CF) [23] and Cisplatin Resistance set (CR), curated genes list from the latest literature with cisplatin resistance. Rectangles with star * show significant enrichment pathways of reference gene sets (p-adj values < 0.05).

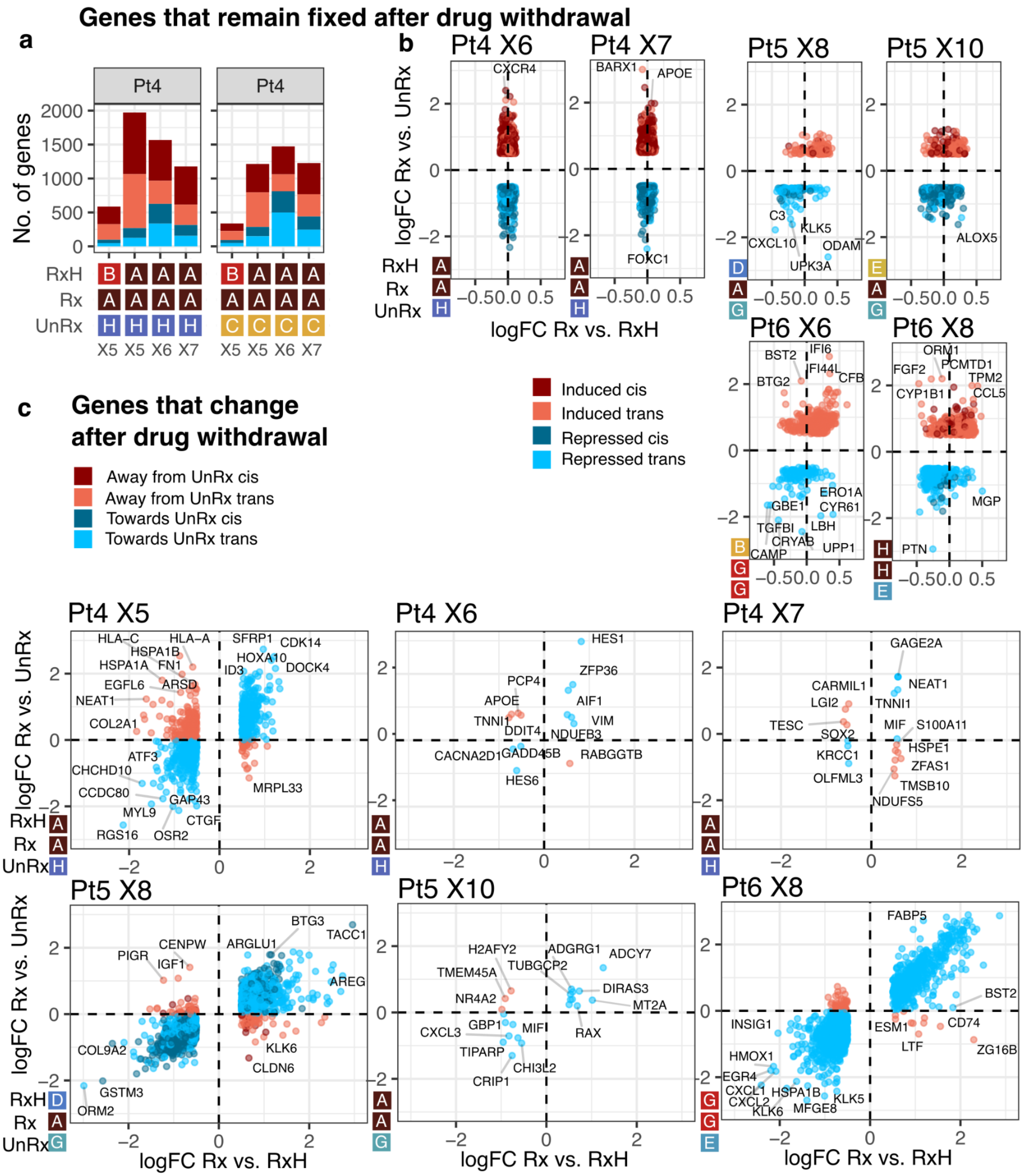


**Supplementary Figure 9: Gene level dynamic changes, supplementary panels to Figure 4.**

(a,b) Genes that remain fixed after drug withdrawal, selected by intersecting the non-differentially expressed Rx vs. RxH genes (FDR > 0.1) with the differentially expressed Rx vs. UnRx genes (FDR < 0.01, |log2 fold change| = |logFC| > 0.5), as in Figure 4a,b. (a) Pt4, comparison of Rx and RxH clones against UnRx clone H, same as in Figure 4b left panel, and comparison against UnRx clone C. (b) Scatter plots for logFC of Rx vs. UnRx (y axis) against logFC of Rx vs. UnRx (x axis), for the selected clones, see also Figure 4b. Each point is a gene. (c ) Genes that change after drug withdrawal, selected as differentially expressed at Rx vs. RxH (FDR < 0.01, |log2 fold change| = |logFC| > 0.5) and intersected with all the genes in Rx and UnRx, as in Figure 4a,e. The panels show scatter plots for logFC of Rx vs. UnRx (y axis) against logFC of Rx vs. UnRx (x axis), for the selected clones, see also Figure 4e.

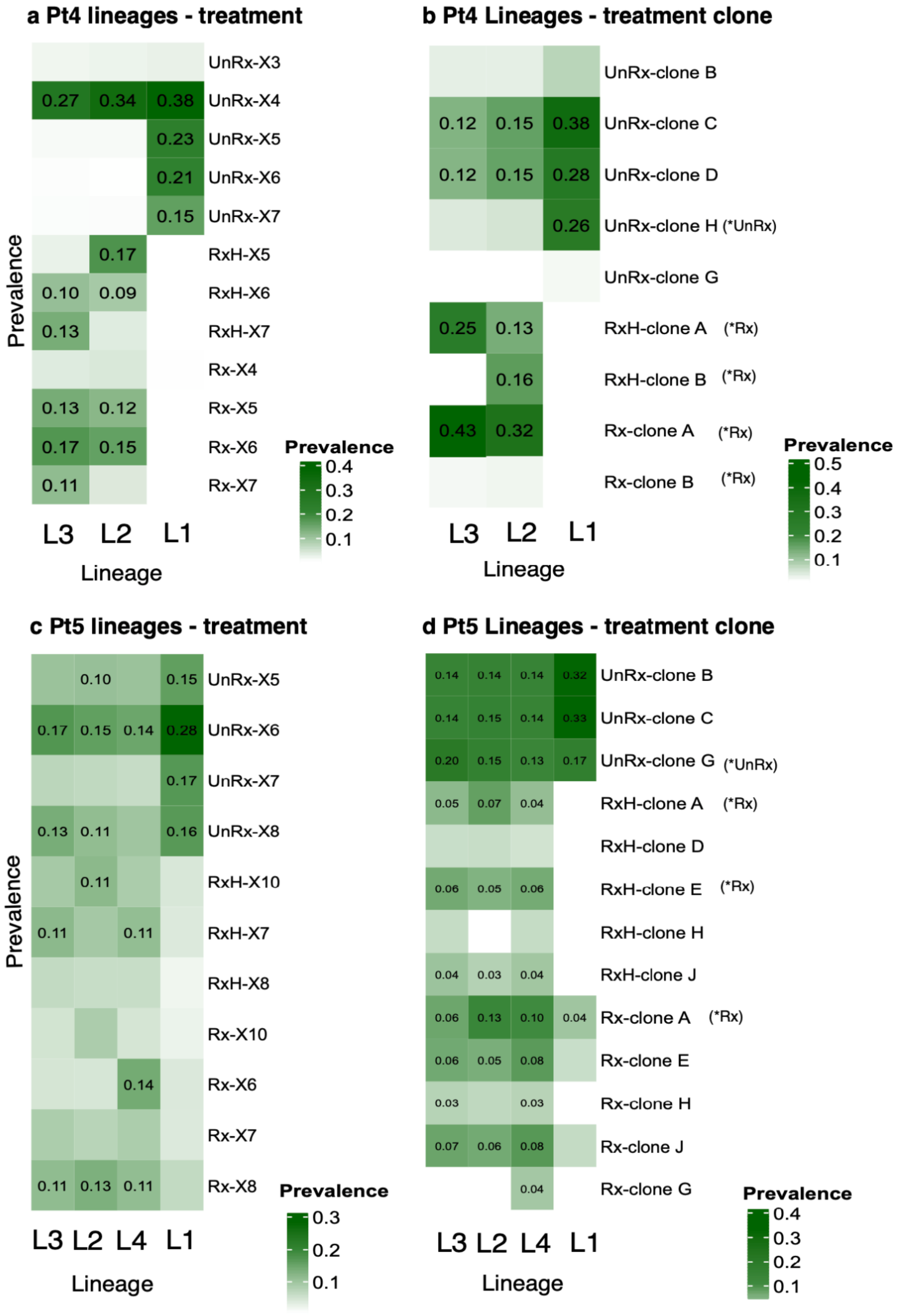
a Pt4 lineages - treatment
UnRx-X3
0.27 0.34 0.38 UnRx-X4
0.23 UnRx-X5
0.21 UnRx-X6
0.15 UnRx-X7
0.17 RxH-X5
0.10 0.09 RxH-X6
0.13 RxH-X7
Rx-X4
0.13 0.12 Rx-X5
0.17 0.15 Rx-X6
0.11 Rx-X7
Prevalence
L3 L2 L1
Lineage
Prevalence
0.4
0.3
0.2
0.1
b Pt4 Lineages - treatment clone
UnRx-clone B
0.12 0.15 0.38 UnRx-clone C
0.12 0.15 0.28 UnRx-clone D
0.26 UnRx-clone H (*UnRx)
UnRx-clone G
0.25 0.13 RxH-clone A (*Rx)
0.16 RxH-clone B (*Rx)
0.43 0.32 Rx-clone A (*Rx)
Rx-clone B (*Rx)
L3 L2 L1
Lineage
Prevalence
0.5
0.4
0.3
0.2
0.1
c Pt5 lineages - treatment
0.10 0.15 UnRx-X5
0.17 0.15 0.14 0.28 UnRx-X6
0.17 UnRx-X7
0.13 0.11 0.16 UnRx-X8
0.11 RxH-X10
0.11 0.11 RxH-X7
RxH-X8
Rx-X10
0.14 Rx-X6
Rx-X7
0.11 0.13 0.11 Rx-X8
Prevalence
L3 L2 L4 L1
Lineage
Prevalence
0.3
0.2
0.1
d Pt5 Lineages - treatment clone
0.14 0.14 0.14 0.32 UnRx-clone B
0.14 0.15 0.14 0.33 UnRx-clone C
0.20 0.15 0.13 0.17 UnRx-clone G (*UnRx)
0.05 0.07 0.04 RxH-clone A (*Rx)
RxH-clone D
0.06 0.05 0.06 RxH-clone E (*Rx)
RxH-clone H
0.04 0.03 0.04 RxH-clone J
0.06 0.13 0.10 0.04 Rx-clone A (*Rx)
0.06 0.05 0.08 Rx-clone E
0.03 0.03 Rx-clone H
0.07 0.06 0.08 Rx-clone J
0.04 Rx-clone G
L3 L2 L4 L1
Lineage
Prevalence
0.4
0.3
0.2
0.1

**Supplementary Figure 10**: **Characterizing each lineage based on the fraction of cells, fitted clone labels, and treatment conditions.** (a) Lineage versus treatment comparison for Pt4. UnRx: untreated, Rx: treated, RxH: drug holiday. (b) Lineage versus clone comparison for Pt4. *UnRx: clone that highly fitted in untreated condition, *Rx: clone fitted under drug, *RxH: clone fitted under drug holiday; In details, lineage L1 - Pt4: high prevalence in untreated cells UnRx; lineage L2-Pt4: large proportion of cells at drug holiday RxH passage X5, X6; lineage L3-Pt4: high prevalence in treated cells Rx and drug holiday late time point RxH passage X7 (c) Lineage - treatment conditions, (d) lineage - treatment clone proportion  correspondence for Pt5; In details, lineage L1 - Pt5: high prevalence in untreated cells UnRx; lineage L2-Pt5: large proportion of cells at first time treatment Rx passage X6; lineage L2-Pt5, L3-Pt5: high prevalence in treated cells Rx and drug holiday RxH.

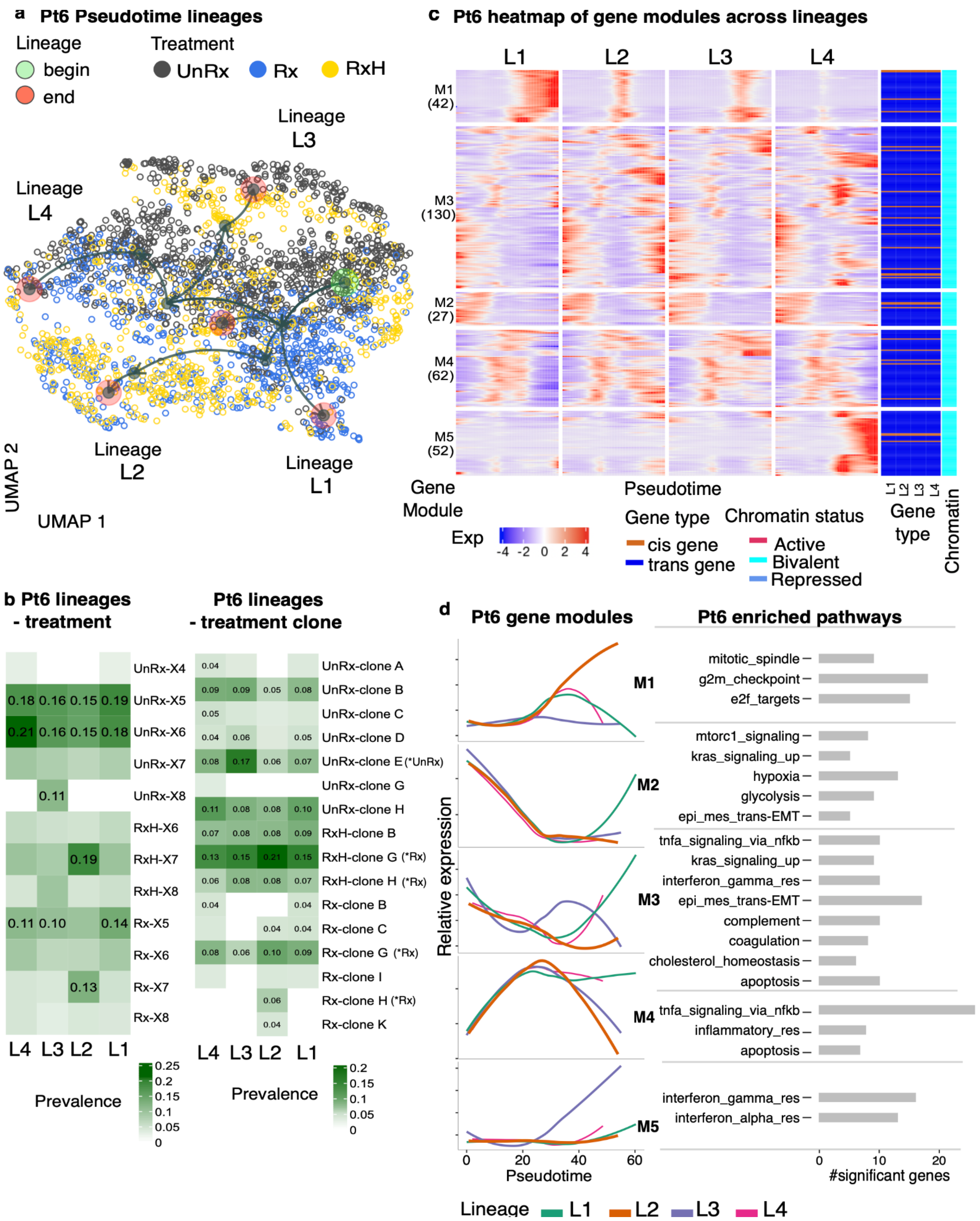

a Pt6 Pseudotime lineages
Lineage
begin
end
Treatment
UnRx
Rx
RxH
Lineage L3
Lineage L4
Lineage L2
Lineage L1
UMAP 2
UMAP 1
c Pt6 heatmap of gene modules across lineages
L1
L2
L3
L4
M1 (42)
M3 (130)
M2 (27)
M4 (62)
M5 (52)
Gene Module
Pseudotime
Exp
-4 -2 0 2 4
Gene type
cis gene
trans gene
Chromatin status
Active
Bivalent
Repressed
Gene type
Chromatin
b Pt6 lineages - treatment
UnRx-X4
UnRx-X5
UnRx-X6
UnRx-X7
UnRx-X8
RxH-X6
RxH-X7
RxH-X8
Rx-X5
Rx-X6
Rx-X7
Rx-X8
0.18 0.16 0.15 0.19
0.21 0.16 0.15 0.18
0.11
0.19
0.11 0.10 0.14
0.13
L4 L3 L2 L1
Prevalence
0.25 0.2 0.15 0.1 0.05 0
Pt6 lineages - treatment clone
UnRx-clone A
UnRx-clone B
UnRx-clone C
UnRx-clone D
UnRx-clone E (*UnRx)
UnRx-clone G
UnRx-clone H
RxH-clone B
RxH-clone G (*Rx)
RxH-clone H (*Rx)
Rx-clone B
Rx-clone C
Rx-clone G (*Rx)
Rx-clone I
Rx-clone H (*Rx)
Rx-clone K
L4 L3 L2 L1
Prevalence
0.2 0.15 0.1 0.05 0
d Pt6 gene modules
Pt6 enriched pathways
M1
M2
M3
M4
M5
Relative expression
Pseudotime
0 20 40 60
mitotic_spindle
g2m_checkpoint
e2f_targets
mtorc1_signaling
kras_signaling_up
hypoxia
glycolysis
epi_mes_trans-EMT
tnfa_signaling_via_nfkb
kras_signaling_up
interferon_gamma_res
epi_mes_trans-EMT
complement
coagulation
cholesterol_homeostasis
apoptosis
tnfa_signaling_via_nfkb
inflammatory_res
apoptosis
interferon_gamma_res
interferon_alpha_res
0 10 20
#significant genes
Lineage L1 L2 L3 L4

**Supplementary Figure 11: Pseudotime analysis of Pt6 showing dynamic gene regulation across treatment, drug holiday, and untreated time points.**

(a)UMAP visualization of cells in all lineages - output of pseudotime analysis coloured by actual drug treatment status (UnRx: untreated cells, Rx: drug treatment, RxH: drug holiday, and time point X (i.e. X4, X5,...). Individual lineage with its starting point - green, and end point - red circles.

(b) Characterization of pseudotime lineages based on cell prevalences from actual drug treatment status and from clone labels assigned to cells in each lineage. Clones with high fitness coefficient are noted (*UnRx, *Rx).

(c ) Heatmap of gene expression across different lineages. X-axis: smoothed gene expression in each lineage, color denoting gene expression level, Y-axis: individual gene and grouped by regulatory genes modules (M is gene module) and number of genes in each module, ex: M1(42). Right: gene types of each individual gene across different lineages, orange: in cis, darkgreen: in trans. Chromatin status of each gene module: active - red, bivalent-cyan, repressed-blue.

(d) Summary of relative gene expression for individual lineages in each gene module from heatmap panel c and list of hallmark enriched pathways related to each gene module based on enrichment analysis gprofilers statistical tests with P-adj<0.05, the size of bar denotes number of gene in each gene module that belong to significant hallmark genes set.
